# Widespread SARS-CoV-2 brain infection after intranasal challenge leads to innate and anti-viral immune response and interference with neuronal function in K18-hACE2 mice

**DOI:** 10.64898/2026.11.19.767910

**Authors:** Simon De Neck, Rebekah Penrice-Randal, Leandro Xavier Neves, Parul Sharma, Barbara Helminger, Adam Kirby, Daniele Mega, Maximilian Erdmann, Martina Zanella, Annika Jagels, Carla Ruiz-Casas, Francesco Prisco, Edward Emmott, Alaa Othman, Udo Hetzel, James P. Stewart, Anja Kipar

## Abstract

Severe Acute Respiratory Syndrome Coronavirus 2 (SARS-CoV-2), the causative agent of the coronavirus disease 2019 (COVID-19) pandemic, often spreads to the brain in K18-hACE2 transgenic mice after intranasal challenge, leading to widespread neuronal infection. Despite frequent reports of this phenomenon, the effects of SARS-CoV-2 infection on neurons have not yet been fully investigated. Therefore, using the K18-hACE2 mouse model, we performed an in-depth *in situ* characterisation of brain changes associated with SARS-CoV-2 infection, with a specific focus on infected neurons and glial cells. We challenged mice intranasally with the SARS-CoV-2 Delta variant at a low dose of 10^2^ PFU/mouse and investigated the brains 6 or 7 days later with a combination of morphologic (light microscopy, transmission electron microscopy), virologic (RT-qPCR), and bulk multi-omics (transcriptome, proteome, metabolome, lipidome) approaches. In our model, SARS-CoV-2 infected only neurons, where it replicated, as confirmed by ultrastructural analysis and supported by multi-omics results. Despite high viral burden, productive SARS-CoV-2 infection of neurons elicited only minimal degenerative changes and rare cell death; however, the transcriptome suggested possible functional alterations. Besides the previously reported mild inflammatory response, neuronal infection initiated an innate immune and antiviral response as well as a mild glial response. In conclusion, the K18-hACE2 model of intranasal SARS-CoV-2 Delta infection allows to study the pathomechanisms associated with viral neuronal infection without overt cell death, contrasting with other neurotropic viruses.

## Introduction

Severe Acute Respiratory Syndrome Coronavirus 2 (SARS-CoV-2), the causative agent of the coronavirus disease 2019 (COVID-19) pandemic, is mostly known for its respiratory manifestations (1, 2). Nonetheless, the nervous system is particularly relevant in COVID-19, as neurological and neuropsychiatric symptoms have been reported in patients with both acute COVID-19 and post-COVID-19 condition (2–6). In hospitalised COVID-19 patients, cerebrovascular disease (ischaemic, haemorrhagic), acute encephalopathy and coma have been reported as the most prevalent neurological manifestations (3, 4, 6). In post-COVID-19 condition, sleep abnormalities, chronic headaches, “brain fog”, impaired memory, mood impairment and pain syndromes are particularly common neurological manifestations (reviewed in (5)). Several mechanisms have been proposed to explain the neurological involvement in COVID-19, including neuroinflammation secondary to systemic inflammation and direct infection of the central nervous system (CNS) by SARS-CoV-2, although the latter is considered unlikely (2, 7, 8). Indeed, while lesions have been reported in the brains of patients and were summarised as “a pattern of cerebrovascular pathology and microglial-predominant inflammation” (9, 10), SARS-CoV-2 was only rarely detected in the CNS and its presence was never confirmed at the ultrastructural level (7, 9, 10).

*In vitro* studies of SARS-CoV-2’s ability to infect neurons show that, unless human angiotensin-converting enzyme 2 (hACE2) is overexpressed, only limited numbers of human induced pluripotent stem cell-derived neurons become infected (11, 12). Indeed, SARS-CoV-2 infection depends on ACE2 expression, and the virus uses the endocytic route to enter the cell; it can then complete its replication cycle, including releasing new virions (11, 12). Similarly, *in vitro* work on neurons from transgenic mice expressing hACE2 under the cytokeratin-18 promoter (K18) found these cells permissive to SARS-CoV-2 (13). *In vivo*, neuronal infection is also achieved in the K18-hACE2 mice (the B6.Cg-Tg(K18-ACE2)2Prlmn/J mouse strain being the most common) ((14–20); for a complete list of references, see Supplemental Table S1). Here, intranasal challenge with SARS-CoV-2 ancestral strains and several previous variants of concern (VOCs), such as Alpha, Beta and Delta, frequently results in widespread neuronal infection (Supplemental Table S1). Based on the infection pattern, the olfactory route is considered the main way of entry to the brain in this model (14, 15, 19, 21– 23). These findings provide clear evidence that SARS-CoV-2 is neurotropic under certain conditions, supporting investigations into its primary routes of entry and the consequences for infected neurons. Interestingly, despite the abundance of studies in K18-hACE2 mice, in-depth approaches to this topic are sparse. Evidence of neuronal damage due to SARS-CoV-2 infection, either at the light microscopic or at ultrastructural level, has been reported but not thoroughly investigated (15, 17, 19, 23–28). Additionally, while it is agreed that SARS-CoV-2 infects neurons in the brain of K18-hACE2 mice, some publications also suggest infection of astrocytes, microglia, oligodendrocytes, choroid plexus epithelium and endothelial cells (16, 24, 25, 27, 29). Previous studies by our group found no evidence of glial or endothelial cell infection (14, 30) and no *in situ* evidence of neuronal degeneration or necrosis, even with widespread infection and mild inflammatory infiltration, which is why we became interested in determining which brain cells the virus targets and what effect it has.

This study aimed to confirm that neurons mediate SARS-CoV-2 infection *in vivo* and to determine the morphological and functional effects on neurons. We used the K18-hACE2 mouse model and a highly pathogenic former VOC, SARS-CoV-2 Delta, as a worst-case scenario and combined light microscopy, transmission electron microscopy, virologic and multi-omics (transcriptomic, proteomic, metabolomic, lipidomic) approaches. The results confirm that SARS-CoV-2 can replicate in neurons. This is associated with an innate immune response and evidence of functional neuronal alterations, occasionally resulting in neuronal death, highlighting SARS-CoV-2 as a primarily neurotropic virus that hijacks the cellular machinery.

## Material and Methods

### Virus

The SARS-CoV-2 Delta isolate (B.1.617.2 hCoV-19/England/SHEF-10E8F3B/2021; GISAID accession number EPI_ISL_1731019), an isolate that has been shown to spread to the brain with high frequency after intranasal challenge of K18-hACE2 mice (14, 31), was used for experimental infections.

### Animals and virus infection

Animal work was approved by the local University of Liverpool Animal Welfare and Ethical Review Body and performed under UK Home Office Project Licence PP4715265. It was undertaken in accordance with locally approved risk assessments and standard operating procedures. The study was undertaken on mice carrying the human ACE2 gene under the control of the keratin 18 promoter (B6.Cg-Tg(K18-ACE2)2Prlmn/J) purchased from Charles River Laboratories (Wilmington, Massachusetts, USA). Mice were maintained in biosafety level 3 facilities under SPF barrier conditions in individually ventilated cages.

The study was undertaken on cohorts of female K18-hACE2 mice, aged 6-8 weeks, that were also investigated as part of another study (30). For infections, mice were anaesthetised lightly with isoflurane and inoculated intranasally with 50 µl sterile PBS containing the virus isolate at 10^2^ plaque-forming units (PFU). Mock-infected mice, injected intranasally with 50 µl sterile PBS, served as controls. The mice were monitored daily for clinical signs, weighed, and euthanised by an overdose of pentobarbitone after reaching the clinical endpoint (maximum of 20% loss of body weight), or at the end of the experiment, at 6 or 7 days post-infection (dpi) (Supplemental Table S2). Mice were dissected immediately after death, and the brains were collected and sampled for downstream processing as part of the general sampling protocol (30).

### Tissue processing

From all mice, as reported in (30), a slice of brain (approximately 1 mm thickness) was prepared by coronal sections at the level of the hippocampus and the thalamus (one section rostral, the second caudal to the pituitary ring) from which a sample (2-3 mm^3^) from the thalamic region was cut out and fixed in 5% glutaraldehyde (GA) (buffered in 0.2 M cacodylic acid buffer, pH 7.3) for transmission electron microscopy (TEM); further samples were collected for RT-qPCR, bulk transcriptomic (thalamus), bulk metabolomic and bulk lipidomic (hypothalamus) and bulk proteomic (hippocampus and cerebral cortex) (Supplemental Figure S1) and frozen at -80 °C. The remaining brain tissue was fixed in 10% NBF.

### Histological and immunohistological examinations

All NBF-fixed tissue samples were transferred to 70% ethanol after 24-48 h fixation until further processing. The brain was trimmed by coronal sections, which were then routinely paraffin-wax embedded (30). Consecutive sections (3-4 µm) were prepared and stained with haematoxylin and eosin (HE) or subjected to immunohistochemistry (IH) for SARS-CoV-2 nucleocapsid protein (NP), ionized calcium-binding adapter molecule 1 (IBA1; marker of microglia, monocytes and macrophages), and glial fibrillary acidic protein (GFAP; marker of astrocytes), with 3-3’-diaminobenzidine (DAB) as chromogen, as previously described (14).

In addition to the earlier examinations (30), the HE and SARS-CoV-2 NP-stained brain sections were subjected to a more in-depth investigation of the neuronal cell population, with specific focus on viral NP expression patterns and evidence of cell degeneration. The sections stained for SARS-CoV-2 NP, IBA1 and GFAP were examined to assess the DAB reaction pattern and the cell morphology of the GFAP- and IBA1-positive cells, focusing on the brain regions selected for image analysis (see below).

### Image analysis

The expression of SARS-CoV-2 NP, IBA1, and GFAP was quantified in the IH-stained sections with QuPath (v0.7.0) (32), by determining the area of DAB deposition for each stain. The slides were scanned using a NanoZoomer 2.0-HT scanner (Hamamatsu, Hamamatsu City, Japan). Using Allen’s Brain Atlas (33–36), regions of interest (ROIs) were manually drawn to include the brain samples belonging to cerebral nuclei, interbrain, midbrain, and/or hindbrain (Supplemental Figure S1A). Because these four regions were not present in all brain section (a direct consequence of extensive sampling for virologic and multi-omics analyses), we grouped them as cerebral nuclei/interbrain and midbrain/hindbrain to improve cohort comparisons. We further refined the tissue selection using a trained pixel classifier (ANN_MLP) to exclude background, and detected DAB using a pixel threshold with a custom threshold for each marker to maximise DAB detection while limiting false positives. The relative area of DAB deposition for each sample was expressed as the ratio between the DAB area and the brain region area. All statistical analyses were performed in R (v4.5.2 (37)) with RStudio (v2026.01.0 Build 392, Posit, Boston, USA), and the comparison between cohorts was done with a Mann-Whitney U test.

For IBA1, a second analysis focused on characterising IBA1-positive detections (referred to as IBA1-positive cells from now on). Using the same ROIs as for assessing IBA1 relative area, we segmented the cells of interest and computed several measurements. First, using the same pixel classifiers as for the IBA1 relative area approach, the IBA1-positive cells were segmented using QuPath (32). Next, we filtered out small annotations, estimated each cell’s soma area using an erosion-dilation step, excluded cells without a soma, and, if multiple soma were detected in a cell, kept only the largest one. These steps aimed to retain only IBA1 detections that could qualify as cells and to exclude detections corresponding only to segments of cell processes. Additionally, we computed the convex hull of each cell and the distance between the closest soma, cell, and hull. From each IBA1-positive cell, a binary mask was exported from QuPath and was used to perform the skeleton, fractal, and Sholl analysis in Fiji (v2.18.0/1.54p (38)). For the skeleton analysis, Skeletonize (2D/3D) (version 2.1.1) and Analyze Skeleton (2D/3D) (version 3.4.2) were used (39). The fractal analysis was performed with the FracLac plugin (version 2016Apr120248a502) (40), following previously published settings (41). Finally, a Sholl analysis (42) (using the SNT plugin version 5.0.14 (43)), and a skeleton analysis (using “Skeletonize” and “Analyze Skeleton (2D/3D)”) focusing only on the cell processes of each IBA1-positive cell, was conducted using a modified version of a published script (44). Additionally, several measurements were computed based on the results from QuPath, skeleton, and Sholl analysis. Details on the image analysis pipeline, as well as the list of all measurements performed on the IBA1-positive cells, are available in the Supplemental Material and Methods. All measurements were imported into R/RStudio for the data analysis. Briefly, the results were investigated at the sample level (median of each measurement) and at the cell level (clustering analysis) (45). The results of the clustering analysis were exported to QuPath for visual assessment of the clusters. Details on the data analysis pipeline are provided in Supplemental Material and Methods.

### Transmission electron microscopy (TEM) including immune-EM

The GA-fixed brain specimens were routinely embedded in epoxy resin, sectioned, processed, contrasted, immunostained for SARS-CoV-2 NP (for immune-EM) and viewed as previously described (30). Sections of an infected hACE2-transfected HEK-293T cell pellet (30) served as positive controls to assess the ultrastructural morphology of SARS-CoV-2 (46–52) and to validate the immune-EM against SARS-CoV-2 NP. Sections from the pellet and brains incubated without the primary antibody, as well as the uninfected brains, served as negative controls. Ultrastructural pictures from the brain of both SARS-CoV-2 Delta-infected and mock-infected cohorts were assessed for evidence of morphological changes, with a particular focus on neurons (53). Specific emphasis was laid on identification of SARS-CoV-2 target cell types, hallmarks of SARS-CoV-2 replication (such as double-membrane vesicles (DMVs) and viral particles (15)), neuron and glial cell morphology, and changes consistent with cell degeneration/death.

### RNA extraction and quantitative RT-qPCR for viral load

RNA was extracted and SARS-CoV-2 quantification was performed using the nCOV_N1 primer/probe mix from the SARS-CoV-2 (2019-nCoV) CDC qPCR Probe Assay (IDT), as previously described (30, 31).

### Omics analysis

Brain samples were collected and processed for bulk transcriptomic (thalamus), untargeted bulk proteomic (cerebral cortex and hippocampus), metabolomic (hypothalamus), and lipidomic (hypothalamus) analysis (Supplemental Figure S1), as already described (30). The specifics of the analysis for the present study are briefly described below, and in more detail in Supplemental Material and Methods.

### Single-omic analyses

We investigated the transcriptomic and proteomic datasets with a focus on SARS-CoV-2 replication, innate immunity, antiviral response, glial response, and neurons. A focus was also placed on the SARS-CoV-2 proteome. In addition to functional enrichment with gene set enrichment analysis (GSEA) using Gene Ontology (GO) (54, 55), we also performed enrichment analysis against the Reactome database (56–58), using the ReactomePA package with GSEA (59). Results were visualised with volcano plots, principal component analysis (PCA), heatmaps, dot plots, and bar plots (30), as well as enrichment map, using clusterProfiler(); for GO enrichment, results were summarised with the clusterProfiler::simplify() function (60). We considered gene products to be significantly different if the false discovery rate (FDR) was less than 0.05, and the log_2_ fold change (log_2_FC) was greater than |2| (transcriptome) or |0| (proteome).

Using CIBERSORTx (61), we estimated proportions of different cell types in the bulk transcriptome dataset. Briefly, we built a reference matrix of murine thalamus using non-normalised gene expression data (averaged UMI counts per cell type; Peter Lönnerberg, personal communication) (62) and used it as input for deconvolution, together with the transcriptome data from our samples (non-filtered and non-normalised).

For the metabolomic and lipidomic datasets, we complemented our previous analyses (30) with functional enrichment analysis using FELLA (63) and the KEGG database (64–66) for the metabolome, and LSEA for the lipidome (67) with the LION ontology (68, 69). For the lipidome dataset, clustering of the samples was investigated with an Orthogonal Partial Least Squares Discriminant Analysis (O-PLS-DA) (ropls, (70)), based on the respective groups, in addition to PCA.

### Multi-omics integration

Low-level integration of the single-omic datasets was performed using both an unsupervised (Common Dimension method (ComDim)) and a supervised (consensus orthogonal PLS-DA (O-PLS-DA) (71)) approach. We visualised and explored the results using block contributions, scores and loadings plots, as well as variable influence on projection (VIP) versus loadings on the predictive component. Additionally, we performed high-level integration of the transcriptome and proteome by comparing both gene products and functional enrichment results. Details on the methods are available in Supplemental Material and Methods.

## Results

### SARS-CoV-2 Delta infects and replicates in neurons in the brain of K18-hACE2 mice after intranasal challenge

#### *In situ* evidence of viral replication and associated cellular changes

Following our previous findings that intranasal challenge of K18-hACE2 mice with the SARS-CoV-2 Delta variant at 10^2^ PFU/mouse leads to widespread neuronal infection (30) and is associated with a very mild nonsuppurative encephalitis, we conducted further in-depth investigations of the brains, with a specific emphasis on neurons. Confirming what we and others have reported previously for several SARS-CoV-2 isolates (for a full list of references, see Supplemental Table S1), SARS-CoV-2 was exclusively detected in neurons; we confirmed this based on the presence of viral NP and/or viral structures identified at the ultrastructural level, and the morphology of the infected cells. Widespread neuronal infection was demonstrated by IH for SARS-CoV-2 NP, as illustrated in (30), showing NP expression in both the cytoplasm and cell processes (i.e. dendrites and axons), sparing the nucleus. The DAB reaction varied in intensity and had a granular or coarse pattern. NP-positive cells often exhibited well-circumscribed small vacuoles and clear cytoplasmic spaces (Figure 1); otherwise, they appeared morphologically unaltered. At ultrastructural level, infection of neurons was confirmed by identification of key elements of the viral replication cycle, such as double-membrane vesicles (DMVs) (Figure 1A, B, D; Figure 2A, B; Supplemental Figure S2), aggregates of NP in the cytoplasm (Figure 1C, Figure 2C), membrane-bound viral particles (compatible with distended endoplasmic reticulum (ER)/ER-to-Golgi intermediate compartment (ERGIC)/lumen of the nuclear envelope and convoluted membranes (CMs)/large virion-containing vacuole (LVCV) (51, 52)) (Figure 1A-D, Figure 2B, C) and virions in the extracellular space (suggestive of viral release from the infected neuron) (Figure 2C), combined with viral NP detection (Supplemental Figure S3A, B) (15, 46–50).

**Figure 1.**
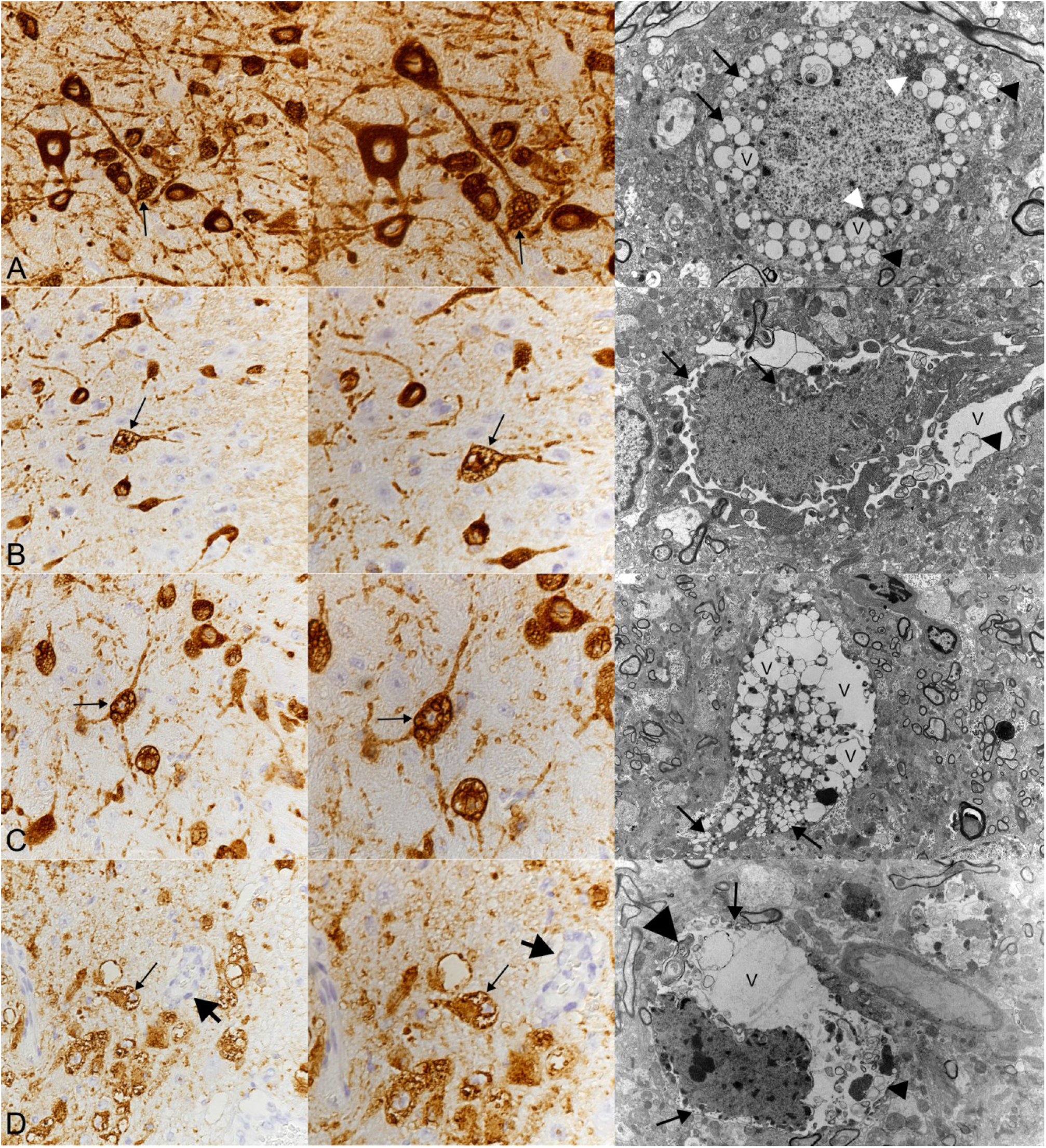
Brain, K18-hACE2 mice after intranasal challenge with SARS-CoV-2 Delta (10^2^ PFU/mouse); 6-7 dpi. Morphological features of infected neurons in brains with widespread neuronal infection, indicated at the light-microscopic level (left column: immunohistochemistry (IH) for SARS-CoV-2 N protein (NP); middle column: higher magnification of the IH), with their likely ultrastructural correlates (right column: transmission electron microscopy (TEM)). **A)** Mouse 1.3 (IH), mouse 1.7 (TEM). The cytoplasm of infected neurons contains several small, round, clear vacuoles (arrows, V). The TEM image shows viral particles in single-membrane vacuoles (arrows), structures consistent with double-membrane vesicles (DVMs; black arrowhead), and aggregates of SARS-CoV-2 NP in the cytoplasm (white arrowhead). **B)** Mouse 1.7 (IH), mouse 1.3 (TEM). The cytoplasm of infected neurons shows clear, irregular vacuoles (arrows, V). The TEM image shows distended cisterna-like spaces separated by small islands of electron-dense cytoplasm, suggestive of distended endoplasmic reticulum. Viral particles (arrows) and a possible DMV (arrowhead) are visible. **C)** Mouse 1.3 (IH), mouse 1.4 (TEM). The cytoplasm of infected neurons contains round, variably sized, clear vacuoles/vacuolation (arrows; V). The TEM image indicates that some of these vacuoles contain viral particles (arrows). **D)** Mouse 1.5 (IH), mouse 1.3 (TEM). The cytoplasm of infected neurons contains large clear vacuoles (arrows; V). The TEM image shows an infected cell with cytoplasm occupied by a large vacuole containing viral particles (arrows), DMVs (small arrowhead), and a multilamellar structure (large arrowhead). Note the mild perivascular infiltrate (short arrows) representing the very mild nonsuppurative encephalitis.

**Figure 2.**
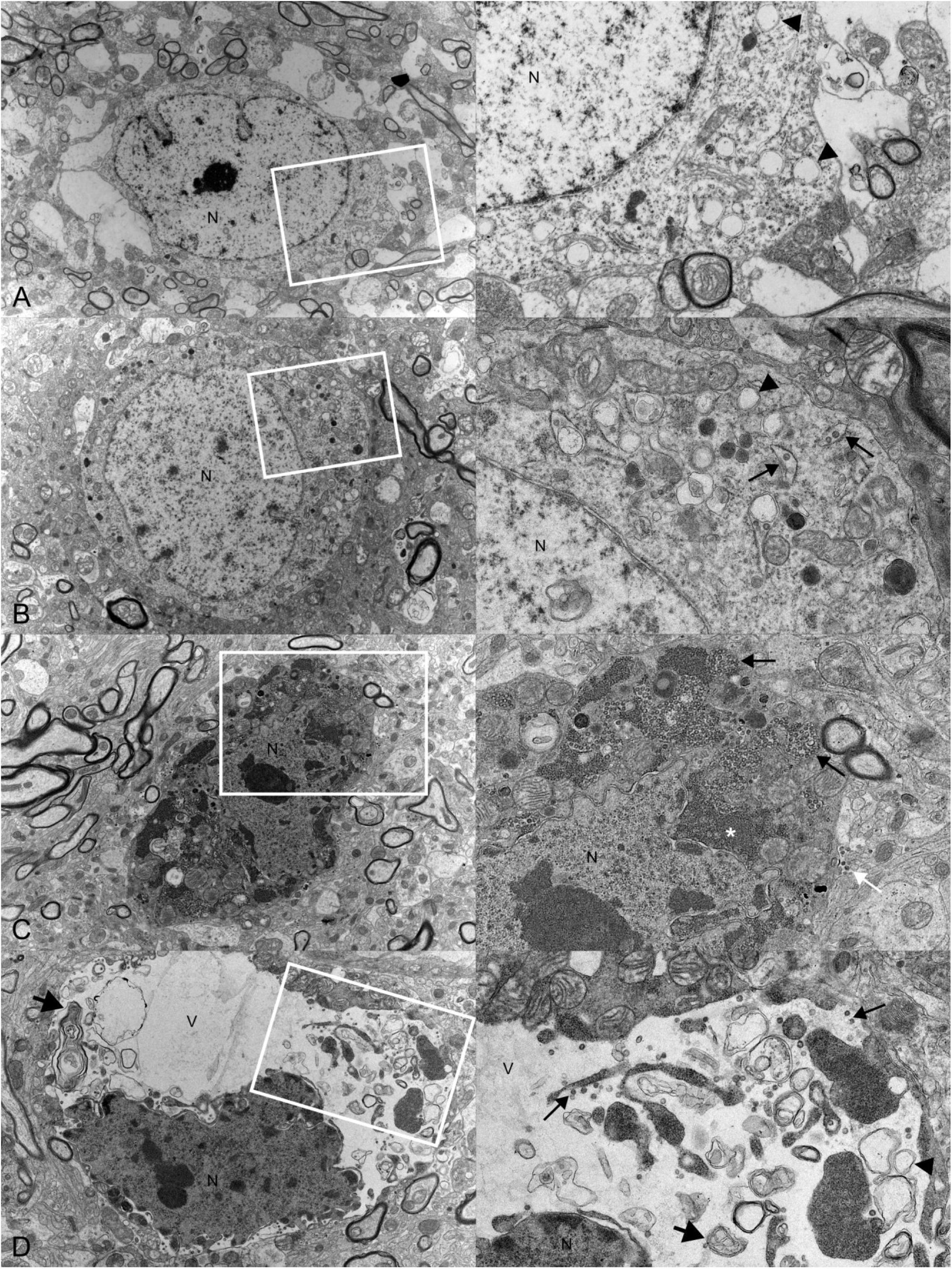
Brainstem, K18-hACE2 mice after intranasal challenge with SARS-CoV-2 Delta (10^2^ PFU/mouse); 6-7 dpi. Illustration of the morphological hallmarks indicating different stages of viral replication in neurons. The pictures on the right are higher magnifications of the region indicated by the white square in the corresponding picture on the left. N: nucleus. **A)** Mouse 1.1. <u>Early viral replication</u>, indicated by occasional double-membrane vesicles (DMVs) (arrowheads) in the cytoplasm. **B)** Mouse 1.7. <u>Advanced viral replication</u>, indicated by more numerous DMVs (arrowheads) and membrane-bound viral particles (arrows) in the cytoplasm. **C)** Mouse 1.3. <u>Late viral replication</u>, indicated by numerous membrane-bound viral particles (black arrows) and large aggregates of SARS-CoV-2 nucleocapsid protein (white asterisk) in the cytoplasm. The nucleus has an irregular contour. Viral particles are present in the extracellular space (white arrow). **D)** Mouse

The ultrastructural replication hallmarks were observed in infected neurons in various combinations; however, based on their presence and abundance, four progressive stages of viral replication and associated neuron “phenotypes” were indicated (Supplemental Table S3; Figure 2; Supplemental Figures S2 and S3) (48). 1) <u>Early viral replication</u>. This was indicated by accumulations of small groups of DMVs in the cytoplasm of otherwise unaltered neurons (Figure 2A). 2) <u>Advanced viral replication</u> was characterised by increased numbers of DMVs, vacuolated structures consistent with distended NE/ER/ERGIC and CMs that contained a few viral particles (membrane-bound viral particles) (Figure 2B) and small aggregates of viral NP in the cytoplasm. 3) <u>Late viral replication</u>. This was indicated by numerous DMVs and abundant membrane-bound viral particles, along with large aggregates of viral NP throughout the cytoplasm or, occasionally, aggregating in the perinuclear region (Figure 2C). The viral particles were contained in single-membrane vacuoles of variable size (compatible with distended NE/ER/ERGIC and CMs/LVCV) (Figure 1A) some of which contained multilamellar structures. In addition, neurons generally showed an irregular nuclear contour (Figure 2C). 4) <u>Exhausting viral replication</u>. This was considered when neurons exhibited distinct morphological alterations, represented by pronounced vacuolation of the cytoplasm, either in the form of distended cisterna-like spaces separated by small islands of electron-dense cytoplasm (suggesting that the latter represent extremely distended ER/ERGIC/nuclear envelope (NE)) (Figure 1B), or more circular to oval vacuolations (suggestive of LVCV) (Figure 1C). Sometimes, one large vacuole almost completely occupied the cytoplasm (Figure 2D). The vacuoles contained variably-sized membranous structures (including DMVs and multilamellar structures) and viral particles. The nucleus had an irregular contour and increased electron density (Figure 2D). 1.3. <u>Exhausting viral replication</u>, indicated by pronounced cytoplasmic vacuolation (V); this is associated with DMVs (arrowhead), viral particles (arrows), and multilamellar structures (short arrows). The nucleus has an irregular contour.

Infected and uninfected neurons were found immediately adjacent to each other (Supplemental Figure S3). Glial cells did not show evidence of viral infection even when observed immediately adjacent to infected neurons (Supplemental Figure S3D).

There was no clear evidence of neuronal death in HE-stained sections. Instead, in both infected and control brains, scattered individual neurons appeared shrunken, with dark basophilic cytoplasm and condensed nucleus (Supplemental Figure S4). Their ultrastructural equivalent was uninfected neurons with an electron-dense, shrunken nucleus with irregular contours, and a shrunken, electron-dense cytoplasm, with distended ER and swollen mitochondria (Supplemental Figure S4). These were interpreted as dark neurons (72, 73) and are considered as the result of cellular contraction, e.g. by post-mortem trauma to an unfixed brain (74).

However, in infected brains, TEM also provided evidence of occasional death of infected neurons. Affected cells exhibited nuclei with large angular electron-dense clumps of chromatin, consistent with pyknosis (Supplemental Figure S5A-C), and occasional widening of the perinuclear space, while the cytoplasm showed features of viral replication. Such degenerate infected neurons were in some instances found with adjacent swollen astrocytic processes (Supplemental Figure S5B). Additionally, we observed phagocytosis of remnants of degenerate cells with structures reminiscent of viral infection, by cells morphologically indicative of microglia and astrocytes (Supplemental Figure S5C, D).

### The brain transcriptome and proteome data support the *in situ* evidence of SARS-CoV-2 replication

The transcriptome and proteomic analyses also supported evidence of viral infection. The proteomic analysis detected nine SARS-CoV-2 proteins, from both the structural (S, N, M), non-structural (pp1ab), and accessory (ORF3a, ORF6, ORF7a, ORF8, ORF9b) categories (Figure 3A, B). All nine proteins were retrieved from five infected brains, while in two brains either ORF6 (mouse 1.8) or both ORF6 and ORF3a (mouse 1.7) were not detected, and in mouse 1.4, only N and ORF9b were detected (Figure 3A).

**Figure 3.**
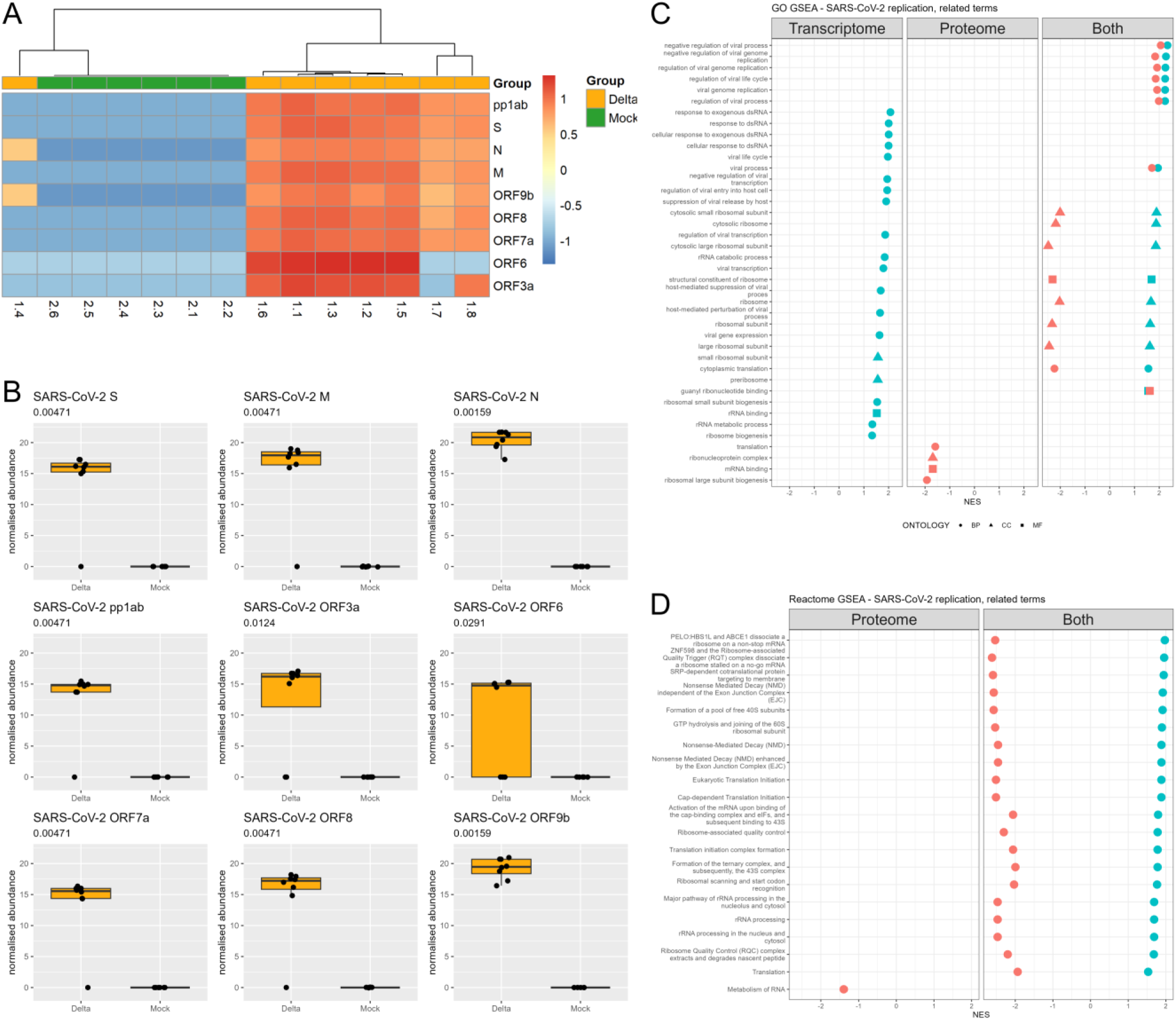
Transcriptome (thalamus) and proteome (cerebral cortex/hippocampus), K18-hACE2 mice after intranasal challenge with SARS-CoV-2 Delta variant (10^2^ PFU/mouse) and mock-infected; 6-7 dpi. A, B) Abundance of SARS-CoV-2 structural (S, M, N), non-structural (pp1ab), and accessory (ORF3a, ORF6, ORF7a, ORF8, ORF9b) proteins. C, D) Functional enrichment analysis using GSEA. **A)** Heatmap illustrating the abundance of SARS-CoV-2 proteins across the samples. The values are (row) scaled for each protein. **B)** Statistical comparison (Mann-Whitney U test) for each viral protein between the Delta-infected and the mock-infected cohort. The p-value of each test is displayed at the top of the corresponding box plot. Illustration of terms from Gene Ontology (GO) (**C**) and Reactome (**D**) which are significantly enriched and related to SARS-CoV-2 replication. BP: biological process, CC: cellular component, MF: molecular function. GSEA: gene set enrichment analysis. NES: normalised enrichment score.

When assessing gene ontology (GO) from the transcriptome and proteome, we observed positive enrichment of terms such as viral life cycle, viral process, viral transcription, viral gene expression, viral genome replication, and their negative regulation (e.g. host-mediated perturbation of viral process), as well as terms associated with double-stranded (ds)RNA and ribosomal (r)RNA (Figure 3C). Additionally, several ribosome-related terms, along with terms associated with mRNA and translation, were either positively or negatively enriched (Figure 3C). For Reactome, terms associated with ribosome, nonsense-mediated decay (NMD), translation, and rRNA were positively or negatively enriched (Figure 3D).

Enrichment analysis of significant metabolites, using the FELLA method (63) with KEGG database (64–66, 75), showed the enrichment of pathways such as “purine metabolism”, “valine, leucine and isoleucine biosynthesis”, and “phenylalanine, tyrosine and tryptophan biosynthesis” (Supplemental Figure S6A). As previously reported (30), neuronal infection with SARS-CoV-2 Delta was not associated with a specific brain lipidome profile, although for 12 lipids (from various classes: LPE, CAR, PC, PS, PI, PE O-) from the positive acquisition mode the abundance differed significantly between the infected and mock-infected cohorts. Unsurprisingly, these lipids were among the most important factors in explaining cohort segregation using O-PLS-DA (Supplemental Figure S7A, B, E). Although no lipids in the negative mode showed significant differences in abundance between the two cohorts, O-PLS-DA clustering was mostly driven by propyl octadecenoic acid, GM3 34:1;O2, tetracosapentaenoic acid, and docosatrienoic acid (Supplementary Figure S7C, D, F). While the two most positively enriched terms in the positive mode were related to TG O- and TG classes (Supplementary Figure S6B), none of the individual lipid species differed significantly in abundance between the cohorts (30), although a cohort-related trend was observed (Supplemental Figure S6C). The multi-omics integration (Supplemental Figure S8-10) showed that several acylcarnitines (such as CAR 16:2 and CAR 18:3, which were significantly more abundant in the infected cohort (30)) and glycerophospholipids with more than two double bonds (polyunsaturated fatty acyl chains: PUFAs; such as PC 36:5, PC 38:6, PC 18:1-18:2, PS 40:8, PS 36:3, PS 44:12, PI 36:5 which were significantly more abundant in the infected cohort (30)), were aligned with gene products related to inflammation/innate immune response (infected cohort) (Supplemental Figure S19D, E), while ether lipids (PC O-34:5, PC O-34:1, PC O-36:2, PE O-16:1_22:4, PE O-40:4; only PE O-40:4 was significantly less abundant in the infected cohort (30)) and cardiolipins (CL 68:6, CL 74:6) were aligned with the mock-infected cohort (Supplemental Figure S19D, F).

### The transcriptome hints at alterations of neuronal function

We used cell-type deconvolution on the bulk transcriptomic data with CIBERSORTx to identify cell-type differences that could inform our understanding of SARS-CoV-2 infection in neuronal cells. Using a custom reference matrix based on a published mouse brain transcriptome (i.e. thalamus), we performed cell deconvolution on the transcriptome (61, 62). For the mock-infected cohort, the results were in line with published molecular murine brain atlases and provided evidence that the proportion of glial cells was slightly higher than the proportion of neurons, and the proportion of excitatory (glutamate) neurons was higher than that of inhibitory (GABA) neurons (62, 76, 77) (Figure 4A, B). Interestingly, in the Delta-infected cohort, the proportions of both excitatory (glutamate) and inhibitory (GABA) neurons were significantly lower, while microglia were higher (Figure 4A, B). However, a subsequent comparison of glutamate and GABA neurotransmitter abundances in the metabolome (available from the hypothalamus) did not detect significant differences between the two cohorts (Figure 4C).

**Figure 4.**
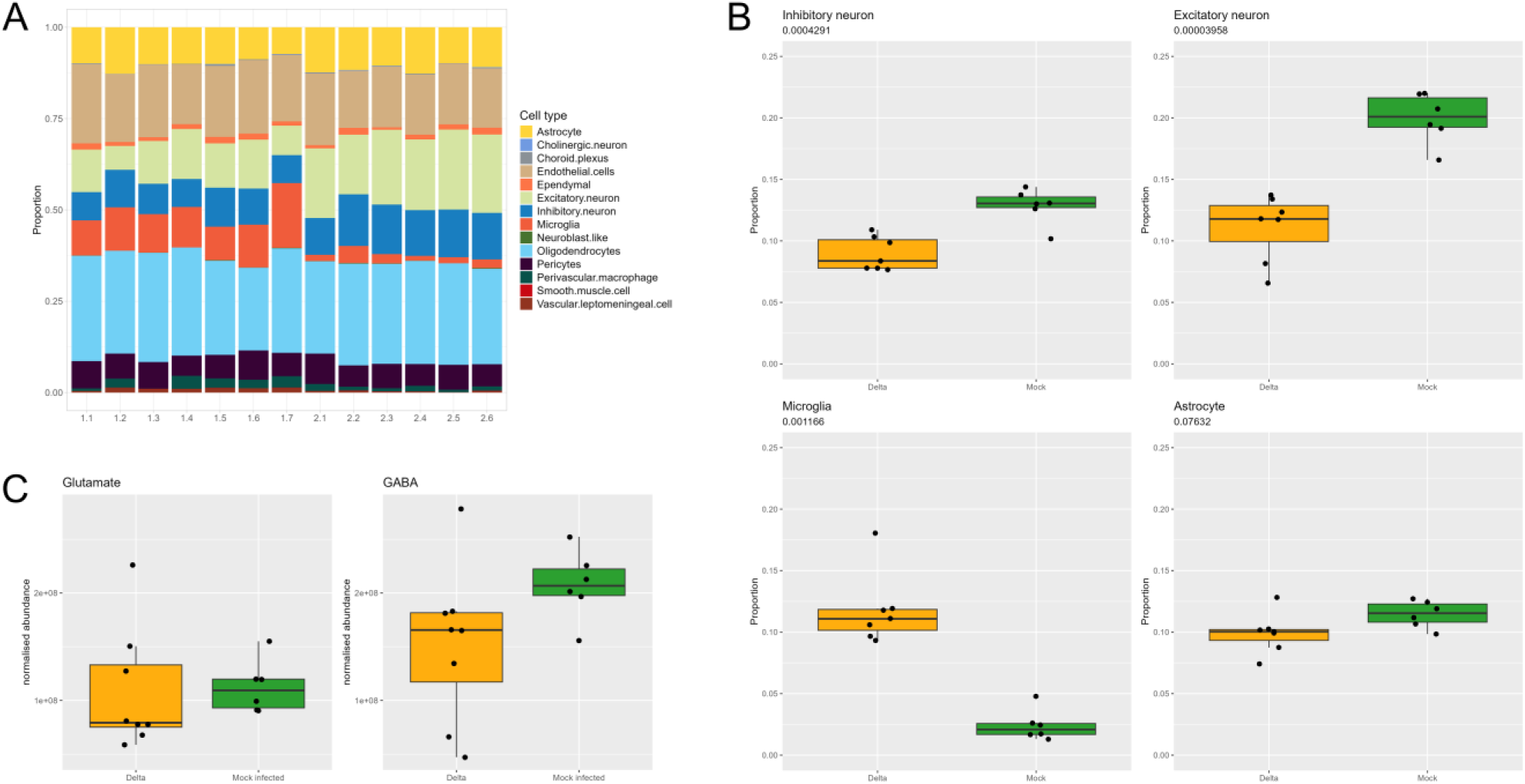
Estimated proportions of cell types based on the brain transcriptome in K18-hACE2 mice after intranasal challenge with the SARS-CoV-2 Delta variant (10^2^ PFU/mouse) and mock-infected; 6 and 7 dpi. **A)** The proportion of the cell types in each sample (bulk transcriptome, thalamus) was estimated with CIBERSORTx (61) using the gene expression in the mouse thalamus as a reference matrix (62). Delta-infected cohort: mice 1.1 - 1.7; Mock-infected cohort: mice 2.1 - 2.6. **B)** Comparison of the estimated proportion of selected cell types determined in the transcriptome (thalamus). Significance was assessed with a t-test (inhibitory neuron, excitatory neuron, astrocyte), or a Mann-Whitney U test (microglia) when the assumptions for a t-test were not fulfilled. The p-value of each test is displayed at the top of the corresponding box plot. **C)** Abundance of the neurotransmitters glutamate and GABA in the metabolome (hypothalamus). No significant difference was observed between the Delta-infected (Delta) and mock-infected cohorts (Mann-Whitney U test).

Transcriptome and proteome parsing for gene products associated with neurons revealed a significant increase in the *Hdc* (histidine decarboxylase; marker of histaminergic neurons (76, 77)) transcript in infected brains (Figure 5A). Three markers of GABAergic neurons (*Gad1, Gad2, Slc32a1*) were slightly but significantly decreased in the transcriptome (76, 77); the difference was not significant in the proteome. Similarly, several neuropeptides (*Pmch, Gcg, Npb, Prok2, Edn1, Oxt, Calcb, Adm, Nmb*), neurotransmitter receptor (*Chrnb1*), and voltage-gated ion channels (*Trpm6, Trpc4*) were significantly up-regulated at the transcript level, while at the protein level only the neuropeptide *Npy1r* was significantly more abundant (Figure 5B) (76–78). In contrast, transcripts of the neuropeptide *Rxfp1* and neurotransmitter receptors (*Oprm1, Glra3*) were detected at significantly lower levels, and the same pattern applied to the neuropeptides *Cck* and *Vgf* in the infected-cohort proteome (76–78). Also, several GABA and glutamate neurotransmitter receptor gene products were significantly less abundant in the infected brains at the transcript level (although only *Gabrb2* reached our threshold of log_2_FC < -2); this was not the case at the protein level (for those gene products that were identified in the proteome) (Figure 5B).

**Figure 5.**
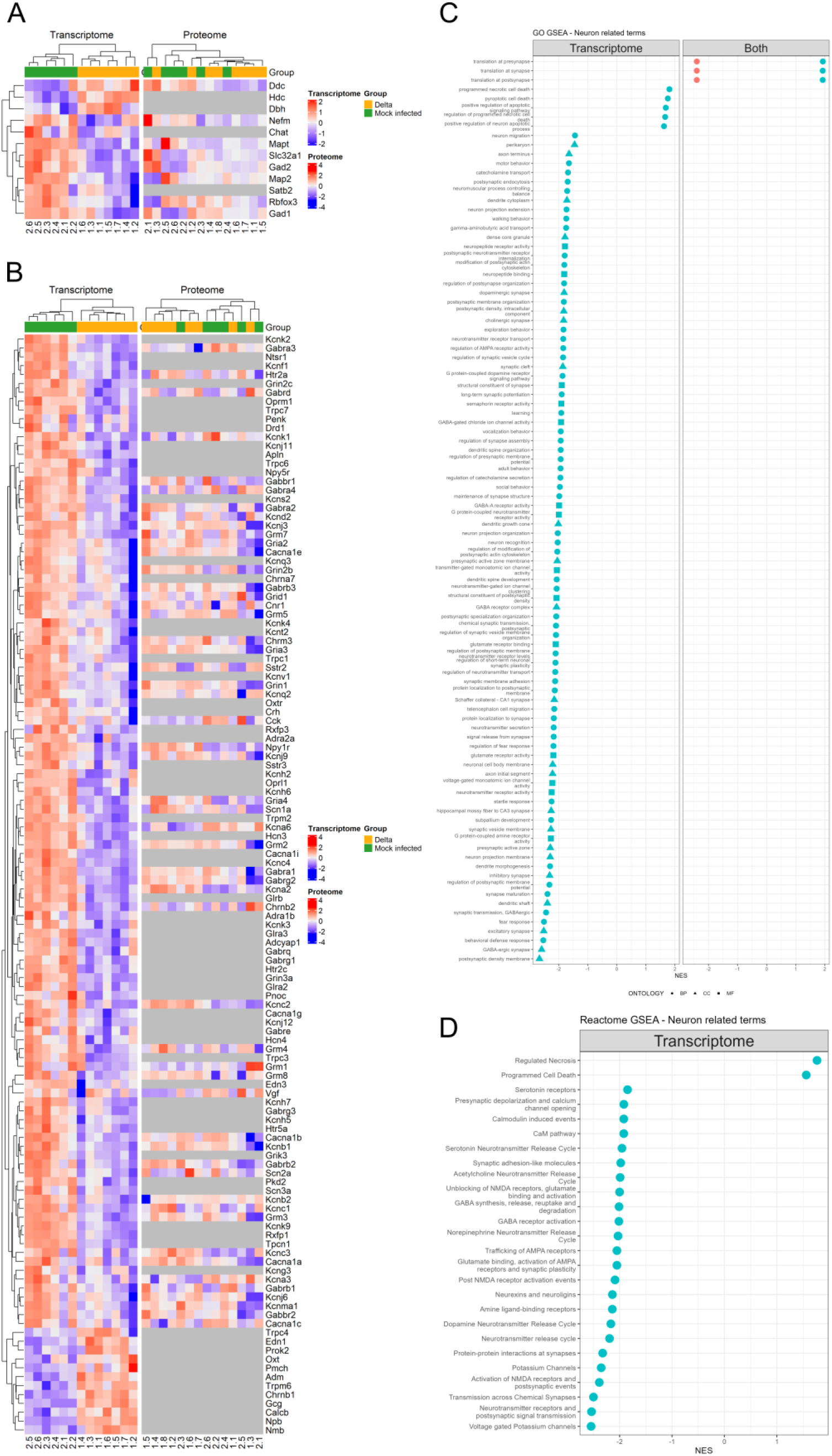
Transcriptome (thalamus) and proteome (cerebral cortex/hippocampus), K18-hACE2 mice after intranasal challenge with SARS-CoV-2 Delta variant (10^2^ PFU/mouse) and mock-infected; 6-7 dpi. **A,B)** Illustration of gene products comparing the Delta-infected (Delta) and mock-infected cohorts for neuronal markers (**A**) and neurotransmitter receptors, neuropeptides, and voltage-gated ion channels (**B**). To compare gene product abundance in the transcriptome and proteome, we mapped significant gene products (False Discovery Rate < 0.05) to significant or non-significant gene products in the proteome and transcriptome, respectively. **C, D)** Functional enrichment analysis using GSEA. Illustration of terms from Gene Ontology (GO) (**C**) and Reactome (**D**) which are significantly enriched and related to neuron morphology and function (e.g. axon, synapse, behaviours), as well as terms related to cell death. BP: biological process, CC: cellular component, MF: molecular function. GSEA: gene set enrichment analysis. NES: normalised enrichment score.

The functional enrichment analysis in the transcriptome and proteome showed positive and negative enrichment of a few terms associated with “translation at synapse” (Figure 5C). Additionally, terms relating to neurons, synapses, and behaviours were negatively enriched in the transcriptome (Figure 5C, D). Of note, several terms associated with cell death (programmed necrotic cell death, pyroptotic cell death, apoptotic signalling pathway, regulated necrosis) were positively enriched in the transcriptome (Figure 5C, D), aligning with the morphological findings.

### SARS-CoV-2 brain infection is associated with the establishment of an innate and antiviral response

We have previously shown that brain infection of the K18-hACE2 mice elicits a mild non-suppurative encephalitis, with leukocyte recruitment and emigration to sites of intense neuronal infection (14, 30). In addition to significant up-regulation of genes (Supplemental Figure S11A, B) and positive enrichment of terms (Supplemental Figure S11C-F) related to inflammation, the top genes and enrichment terms also suggest innate and antiviral responses, such as *Ifit1* and *Ifit2*, or ISG15 antiviral mechanisms (79–81) (Supplemental Figure S11). Top enriched GO terms for the transcriptome further supported the antiviral response, with clusters of terms linked to “regulation of viral life cycle”, “antiviral innate immune response”, and “type II interferon production” (Supplemental Figure S12A, B).

Together, the transcriptome and proteome showed significantly more abundant gene products (i.e. transcripts and proteins) from the innate immune and antiviral response in the infected brains, spanning several categories, such as interferon (*Ifng, Ifnb1*), interferon-stimulated genes (ISG), IFN-inducible GTPases, and interferon regulatory factors (IRF) (Figure 6A, B). Amongst the ISGs, several have been reported to inhibit various steps of the SARS-CoV-2 infection cycle (Figure 6A), such as binding and entry (*Ubd, Clec4d, Ch25h, Cd74, Plscr1*), RNA replication and translation (*Zbp1, Ifit3, Oas2, Oas3*), and assembly and egress (*Bst2, Msr1, Tagap*) (79, 82–86). Three IFN regulators, reported as having a proviral role, were also more abundant (*Isg15, Usp18, Socs1*) (Figure 6A) (83). Additionally, several pattern recognition receptor genes were significantly up-regulated (transcriptome only), including *Tlr1, Tlr2, Tlr3, Tlr7*, and *Nlrp3* (87, 88) (Figure 6C). Functional enrichment analyses (GO and Reactome) aligned with these findings, with positive enrichment of terms related to innate immunity/antiviral response, in both the transcriptome and proteome (Figure 6D, E).

**Figure 6.**
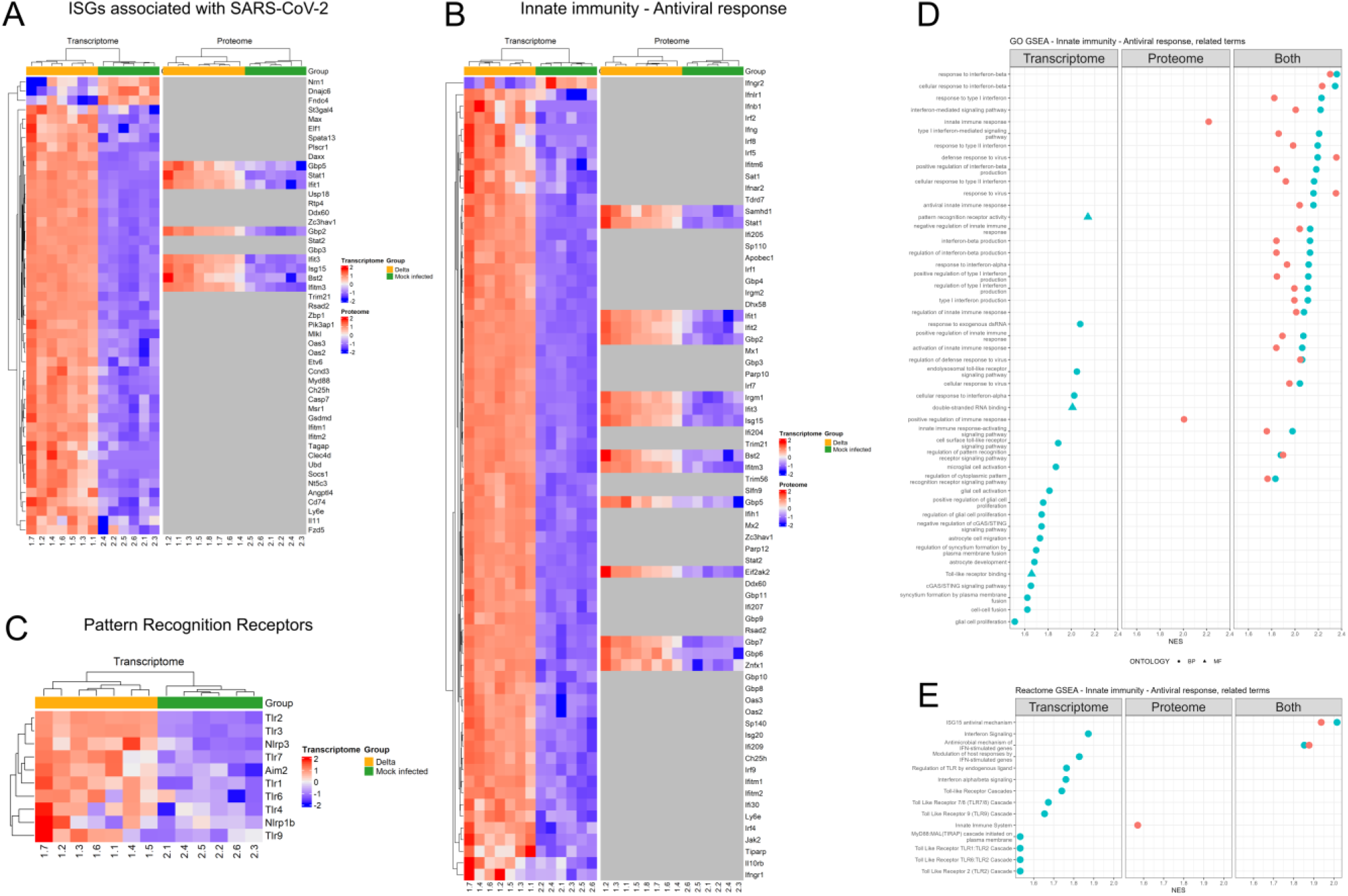
Transcriptome (thalamus) and proteome (cerebral cortex/hippocampus), K18-hACE2 mice after intranasal challenge with SARS-CoV-2 Delta variant (10^2^ PFU/mouse) and mock-infected; 6-7 dpi. **A-C)** Illustration of the significantly different gene products (False Discovery Rate < 0.05) between the Delta-infected (Delta) and the mock-infected cohorts, for: **A)** the interferon-stimulated genes (ISGs) associated with SARS-CoV-2 infection; **B)** the innate immunity and antiviral response; **C)** pattern recognition receptors. **D, E)** Functional enrichment analysis using GSEA. Illustration of significantly enriched Gene Ontology (GO) **(D)** and Reactome (**E**) terms related to innate immunity and antiviral response. BP: biological process, CC: cellular component, MF: molecular function. GSEA: gene set enrichment analysis. NES: normalised enrichment score.

When comparing the transcriptome and proteome in our study to the signature associated with the published inflammation resolution (89), we found gene products from both the induction-selective and resolution-selective phases to be significantly more abundant in the infected cohort (Supplemental Figure S13). However, we noted that the gene products with the highest amplitude of change were from the induction-selective phase. This would align with our morphological findings of different stages of viral replication within infected neurons.

### Neuronal infection by SARS-CoV-2 is associated with a limited, mainly microglial response

Previous studies including our own have reported evidence of a glial cell response in SARS-CoV-2 infected mouse brains (14, 15, 19–21, 26, 27). TEM investigations in the present study showed evidence of occasional phagocytosis of dead cells by microglial cells and astrocytes, consistent with the removal of apoptotic cells (Figure 5C). Hence, we further investigated the role of these glial cells in the observed processes.

### Microglia

The transcriptome-based cell deconvolution suggested a significant increase in the proportion of microglia in the thalamus of the infected cohort (Figure 4A, B). At the same time, a significant up-regulation of microglia markers (*Socs3, Clec7a, Spi1, Csf1, Prprc, Aif1*), including markers of so-called activated microglia (*Itgax, Lgals3, Cd86*) (90–97), was observed, while three markers found in “homeostatic” microglia, *Gpr34, Trem2* and *Fcrl2* (90, 94, 98, 99), were significantly down-regulated (Supplemental Figure S14). The proteome showed no difference for the detected microglia markers (P2Y12 (*P2ry12*), HEXB (*Hexb*), ITAM (*Itgam*)), including AIF-1 (IBA1; *Aif1*), used to identify microglia by IH (90, 94, 97), between the infected and mock-infected cohorts. Several pattern recognition receptors (PRRs) were also significantly up-regulated at the transcript level, either expressed by all glial subtypes, astrocytes, microglia and oligodendrocytes (*Tlr2* and *Tlr3*), or by microglia alone (*Tlr1, Tlr7* and *Nlrp3*) (87, 88). Functional enrichment analysis of the transcriptome showed positive enrichment of GO terms associated with glial cells and microglia specifically, such as “glial cell proliferation/activation” and “microglial cell activation” (Figure 6D). Light microscopy, via IH for IBA1, indicated that the majority of microglial cells in SARS-CoV-2 infected brains exhibited the morphology of reactive microglia (45, 100), as they exhibited a larger soma area and thicker cell processes/branches than the microglia in control brains (Figure 7). Indeed, these prominent microglial cells dominated across the investigated brain regions (cerebral nuclei, interbrain, midbrain, and hindbrain) in most infected brains, mirroring the distribution of infected neurons. A direct association between neuronal infection and microglial reaction was further supported by findings in two mice (1.4, 1.5) with less extensive neuronal infection in the aforementioned regions. Here, only a proportion of microglia displayed the described morphological alterations, and mostly in regions with infected neurons.

**Figure 7.**
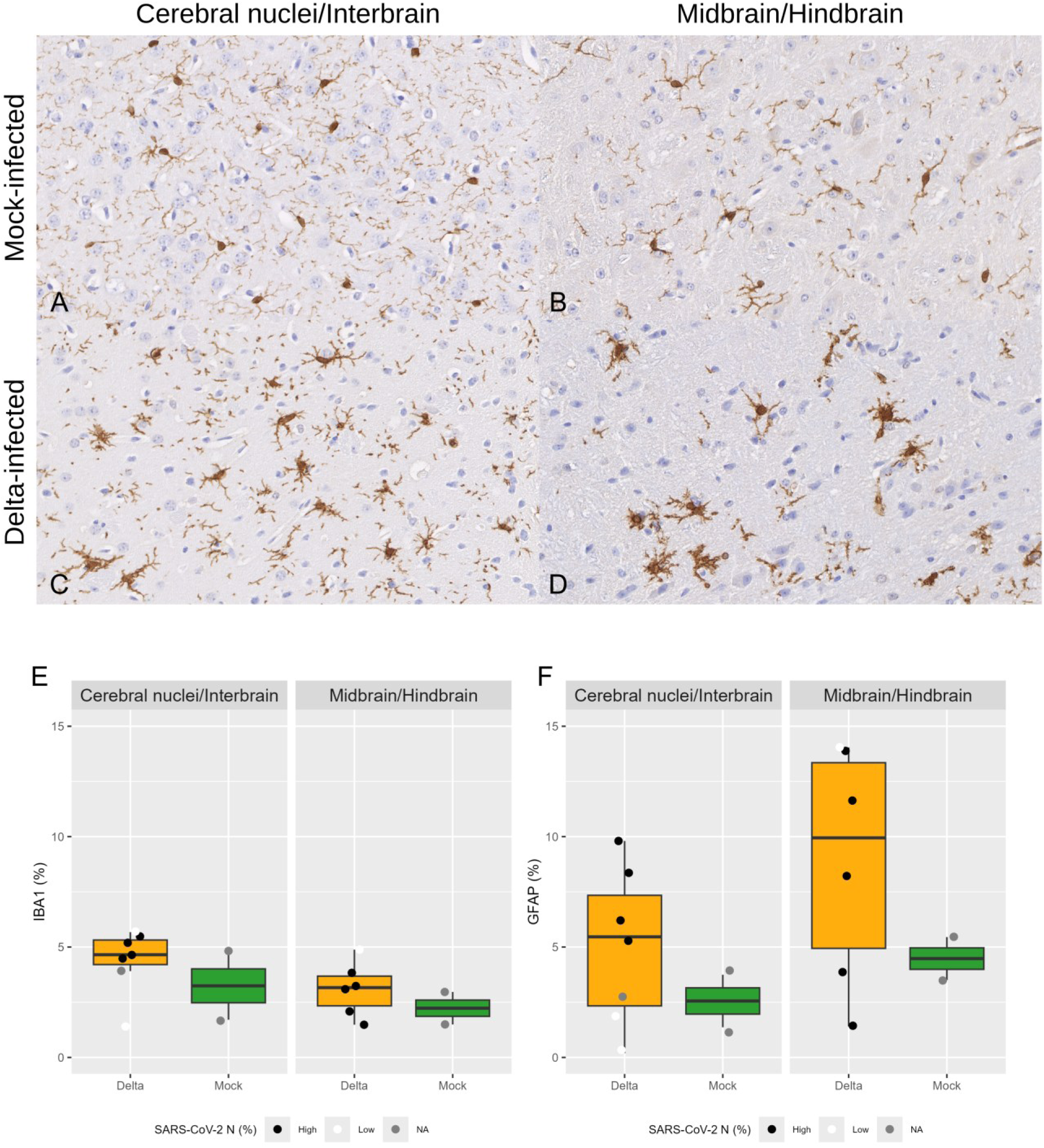
Cerebral nuclei/interbrain and midbrain/hindbrain regions, K18-hACE2 mice after intranasal challenge with SARS-CoV-2 Delta variant (10^2^ PFU/mouse) and mock-infected; 7 dpi. Immunohistochemistry for IBA1 (microglial marker) and GFAP (astrocyte marker). **A-D)** Illustration of microglial shape. **A, B)** Mock-infected cohort, mouse 2.6. Microglial cells have small somas and long thin processes/branches. **C, D)** Delta-infected cohort, mouse 1.7. Microglial cells are prominent due to their larger soma area and thicker cell processes/branches. **E, F)** Quantification of tissue expression of IBA1 and GFAP in the cerebral nuclei/interbrain and midbrain/hindbrain, expressed as the ratio of DAB deposition area to tissue area. Statistical analysis (Mann-Whitney U test) showed no difference between the Delta-infected (Delta) and mock-infected (Mock) cohorts in the proportion of IBA1 (**E**) and GFAP (**F**) immunostaining. The proportion of SARS-CoV-2 NP immunostaining in the corresponding brain region was categorised as either high (> 5% of immunostaining) or low (< 5% of immunostaining). NA: not available.

The image analysis pipeline partially captured altered microglial morphology. While quantification of the total relative area of IBA1 expression in the cerebral nuclei/interbrain and midbrain/hindbrain regions showed no significant difference between the infected and mock-infected cohorts (Figure 7E), cohort-specific differences were visible when assessing the morphological features of individual IBA1-positive cells (Figure 8A, Supplemental Figure 16A-C). At sample levels (i.e. when considering the per-sample median of each morphological measurement), a statistically significant difference between the cohorts (adjusted p-value < 0.05) was found for 21 measurements (Supplemental Table S4) when combining the brain regions, while these were not observed when considering each brain region separately (Supplemental Tables S5, 6). Amongst those, in the infected group, the IBA1-positive cells were more numerous (normalised counts) (Figure 8B, C), a result in line with the cell deconvolution, and had higher tissue density (centroid/border distance between soma, cell, convex hull); they were larger (cell area, soma area, soma perimeter) and had more primary branches (Sholl primary branches) and ramifications (skeleton end-point, Sholl critical value) (Supplemental Table S4). Correlation of the measurements from the image analysis pipeline (i.e. from the cerebral nuclei/interbrain region to match the transcriptome region) with the significantly different genes (FDR < 0.05 and log_2_FC > or < 0) known to be associated with microglia revealed an opposite correlation pattern. As such, measurements positively correlated with up-regulated microglial gene markers were negatively correlated with down-regulated microglial gene markers, and *vice versa* (Supplemental Figure S16). This analysis revealed a positive correlation between up-regulated gene markers and measurements associated with increased cell density, cell and convex hull size, skeleton complexity/ramification, and number of primary branches; there was a negative correlation with the distance between adjacent soma/cell/convex hull. Based on the morphological features, we summarised variation among individual IBA1-positive cells by assigning them to four phenotypes (clusters) (Figure 8D, E; Supplemental Figure S15D-H), the proportions of which were weakly correlated with the cohort (Figure 8F, G). IBA1-positive cells in clusters 1 and 2 were relatively small, with a simple cell skeleton; in clusters 3 and 4 they were larger, with a more complex cell skeleton (Supplemental Figure S15I, J; Supplemental Table S7). The infected cohort tended to have a higher proportion of cluster 3 and 4 cells, and a lower proportion of cluster 1 and 2 cells (Figure 8F, G). An exception was in the cerebral nuclei/interbrain of mouse 1.4, where the cluster pattern was closer to that of the control cohort (Figure 8A, Supplemental Figure S15A). This brain had normalised counts (Figure 8B) and cluster proportions (Figure 8F) in line with the control cohort; it also had the lowest viral load of the cohort, as measured by RT-qPCR (30), and minimal neuronal SARS-CoV-2 NP expression.

**Figure 8.**
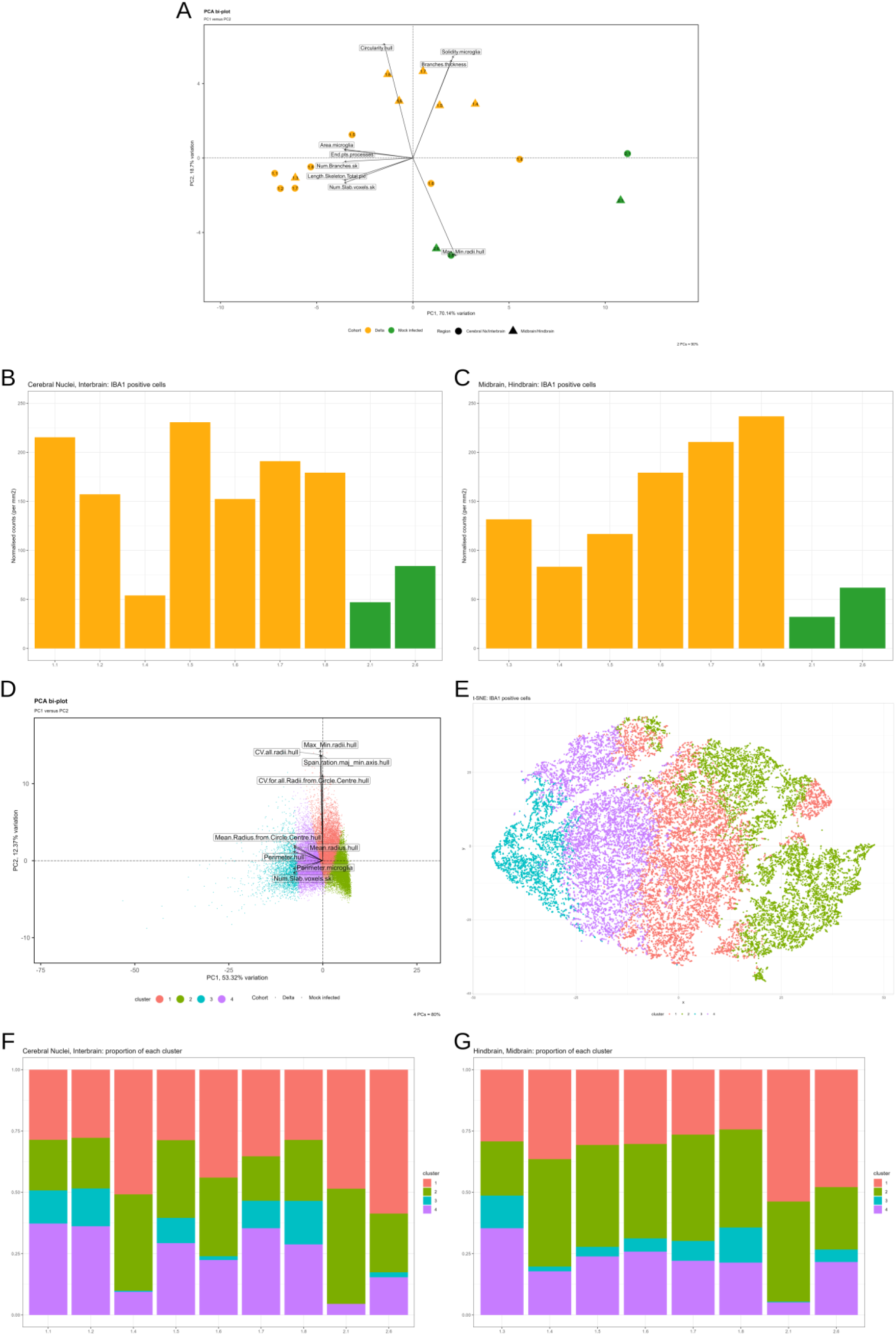
Brain, K18-hACE2 mice after intranasal challenge with SARS-CoV-2 Delta variant (10^2^ PFU/mouse) and mock-infected; 6-7 dpi. Quantification of IBA1 and GFAP expression in the cerebral nuclei/interbrain and the midbrain/hindbrain. **A)** Principal Component Analysis (PCA) showing the clustering of the samples using the median of the measurements derived from the QuPath, Skeleton, Fractal, and Sholl analyses. Samples tend to cluster within their respective groups (i.e., infected and mock-infected). PC: Principal Component. **B, C)** Normalised counts (per mm^2^) of the IBA1-positive cells in the cerebral nuclei/interbrain (**B**) and midbrain/hindbrain (**C**). **D, E)** PCA (**D**) and T-Distributed Stochastic Neighbor Embedding (t-SNE) (**E**) showing the clustering of the individual IBA1-positive cells. The shapes in the PCA biplot correspond to the cohorts. **F, G)** Illustration of the proportion of each of the four clusters in the Delta-infected (1.x) and mock-infected (2.x) cohorts for the cerebral nuclei/interbrain (**F**) and the midbrain/hindbrain (**G**).

### Astrocytes

Although the transcriptome-based cell deconvolution (CIBERSORTx) did not predict a significant difference in the proportion of astrocytes in the thalamus (Figure 4A, B), multiple transcriptomic, proteomic and ultrastructural findings were consistent with astrocyte reactivity as opposed to a change in number. The transcriptome showed significant up-regulation of known astrocyte markers (*Lcn2, Serpina3n, C3, Thbs1*) and reactive astrocyte markers (*Hspb1, Gfap, Stat3, Vim*) (97, 101, 102). Four of these markers were also significantly more abundant at the protein level (SPA3N (*Serpina3n*), NGAL (*Lcn2*), CO3 (*C3*), STAT3 (*Stat3*)) (Supplemental Figure S14), although there was no difference for GFAP (*Gfap*), and FABP7 (*Fabp7*) (astrocyte marker) (97) was significantly less abundant in the infected brains. In addition, the transcriptome showed positive enrichment of the GO terms “astrocyte development” and “astrocyte cell migration” (Figure 6D). Astrocyte morphology, as seen in GFAP-stained sections, showed no obvious differences between infected and mock-infected brains. However, at the ultrastructural level, astrocytes in infected brains exhibited several morphological features of reactive astrocytes: nuclear enlargement, distended cell bodies and processes, and sometimes increased finely granular, electron-dense material in the cytoplasm (indicative of glycogen) (Figure 9). Nonetheless, quantification of GFAP expression in IH-stained sections (expressed as the relative area of DAB deposition) in the cerebral nuclei/interbrain and midbrain/hindbrain regions showed no significant difference between the infected and mock-infected cohorts (Figure 7F, E).

**Figure 9.**
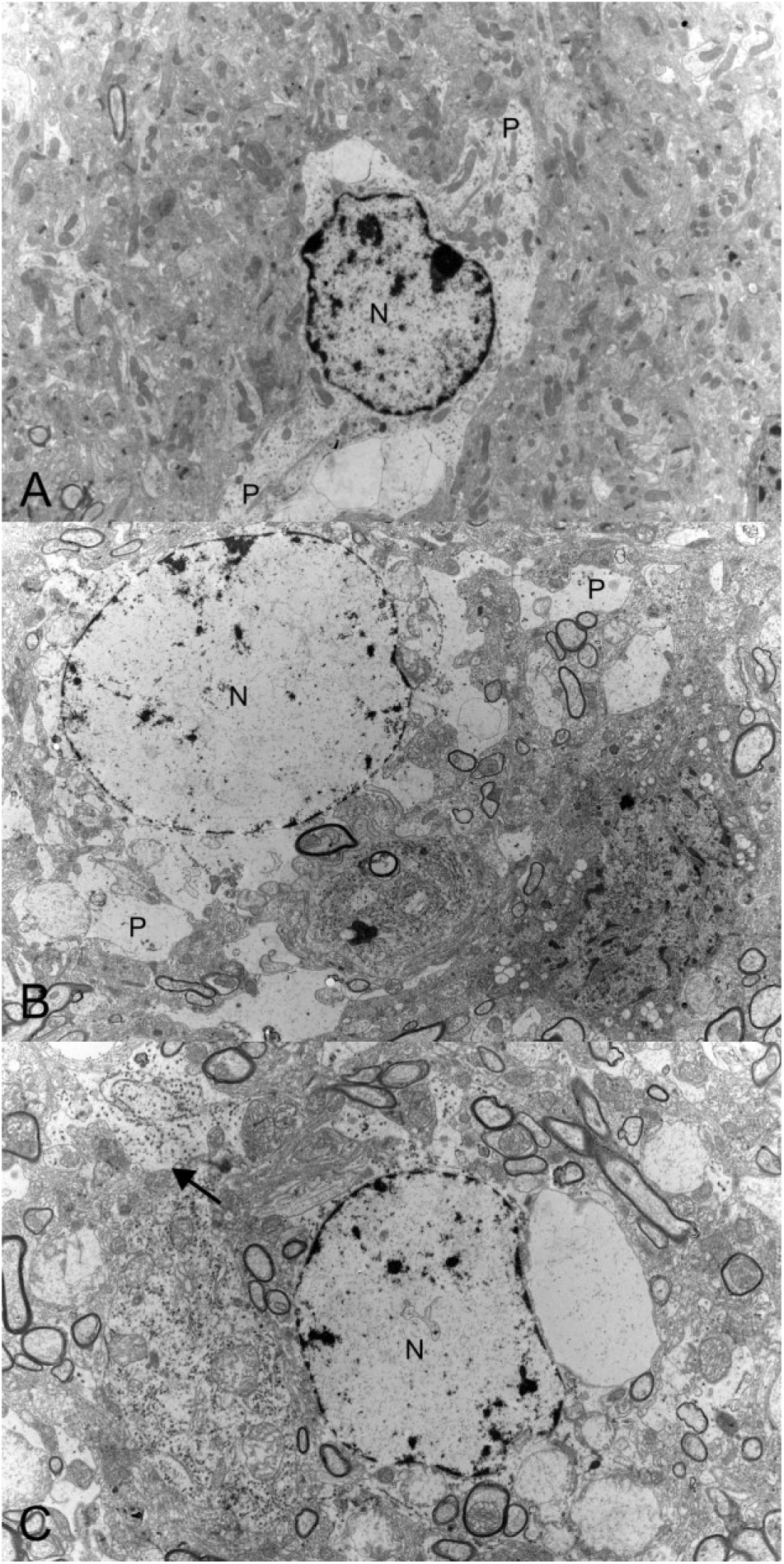
Brainstem, K18-hACE2 mice after intranasal challenge with SARS-CoV-2 Delta variant (10^2^ PFU/mouse) and mock-infected; 6-7 dpi. **A)** Mock-infected mouse (2.1). Astrocyte, with nucleus (N), electrolucent cytoplasm and short processes (P). 4,600x. **B, C)** Delta-infected mice. **B)** Mouse 1.1. Astrocyte with enlarged nucleus (N) and prominent processes (P). The image is taken at the same magnification as A (4,600x). **C)** Mouse 1.1. Astrocytes (N: nucleus) with an increased amount of electron-dense material (compatible with glycogen) in cell processes (arrow). 5,800x.

## Discussion

In this study, combining morphologic and multi-omics approaches, we show that intranasal challenge with SARS-CoV-2 Delta in K18-hACE2 mice leads to viral spread to the brain and replication in infected neurons, where it may induce functional alterations and elicits an innate immune/antiviral response. Productive viral infection in our model is associated with minimal neuronal death, likely due to cellular exhaustion, and a limited glial reaction. These neuron-focused results complement our previous work, which investigated the mechanisms underlying the limited inflammatory response in the brains (30).

We confirmed, in line with previous studies (Supplemental Table S1), that neurons are the SARS-CoV-2 permissive cell type in the brain in K18-hACE2 mice, using complementary *in situ* approaches which detected viral antigen both at light microscopic (IH) and ultrastructural (immune-EM) levels and identified virus replication-associated structures and viral particles (15, 46–52). The accompanying omics datasets supported viral replication in the brain. Proteomics confirmed the presence of structural proteins (S, N, M) as well as non-structural (pp1ab) and accessory (ORF3a, ORF6, ORF7a, ORF8, ORF9b) proteins, which are translated upon viral entry into the permissive cell (103, 104). In addition to viral detection by IH, ultrastructural analysis (immune-EM), and proteomics, transcriptomic and proteomic analyses of the host provided further evidence of viral replication. Functional enrichment analysis showed positive enrichment of terms related to viral genome replication, the viral life cycle, viral process, viral transcription, viral entry, and viral release. These findings support the idea that the presence of the virus alters host transcriptomic and proteomic responses, with effects extending beyond viral entry.

Ribosome-related terms were positively enriched in the transcriptome but negatively enriched in the proteome, while pathways associated with nonsense-mediated decay (NMD) showed a similar divergence. Such discrepancies may reflect SARS-CoV-2-mediated interference with host translation and mRNA processing (104–106), whereby transcriptional responses are not necessarily translated into proportional changes at the protein level. Therefore, the transcriptomic and proteomic datasets may capture different aspects of the host response to infection rather than contradictory biological processes.

To provide additional depth to our analysis, we also performed metabolomic analysis, based on the hypothesis that SARS-CoV-2 replication changes the cellular metabolism. Metabolomics enrichment analysis highlighted KEGG pathways corresponding to “purine metabolism” and “valine, leucine and isoleucine biosynthesis”. Based on *in vitro* work, it is suspected that purine metabolism, and more particularly *de novo* purine synthesis, is a critical step for RNA transcription and the synthesis of subgenomic mRNA (sg-mRNA) in the early phase of the SARS-CoV-2 replication cycle (107). Although the authors of that study reported increased abundance of several purine synthesis intermediates in infected cells, we did not detect these in our metabolomic dataset. In our cohort, the purine metabolism pathway was directly linked to urate, which was significantly more abundant in the infected brains (30). Urate is a product of purine degradation, produced from xanthine by xanthine oxidoreductase (xanthine dehydrogenase/xanthine oxidase), and is further transformed to allantoin by uricase (108, 109). Given that we had performed transcriptomics and proteomics, we interrogated our dataset for xanthine dehydrogenase (*Xdh*) and uricase (*Uox*). *Xdh* was significantly increased in infected brains at both the transcript and protein levels, while *Uox* was significantly decreased at the transcript level and not detected at the protein level. This suggests more intense purine catabolism in infected brains, although we cannot exclude that the increase in urate is part of a protective response. Indeed, urate has antioxidant properties, and an *in vitro* murine model of Parkinson’s disease has shown that increased intracellular urate protects neurons (108, 110). For the “valine, leucine and isoleucine biosynthesis” pathway, a computational modelling study predicted that SARS-CoV-2 depends on isoleucine for replication (111). The metabolome in our study showed no significant differences for valine, leucine, or isoleucine between the Delta-infected and mock-infected cohorts. However, FELLA analysis linked this pathway to 2-methylmaleate (citraconic acid), an immunomodulator with anti-inflammatory, antioxidant, and antiviral functions (112, 113), which was significantly less abundant in infected brains (30).

The lipidome during SARS-CoV-2 infection has been shown *in vitro* to be dysregulated (114); therefore, we explored lipidome changes in our model. Some revealed dysregulation of numerous lipid species, with accumulation of lipids with PUFAs and a reduction in lipid species with saturated and monosaturated fatty acid chains (reviewed in (114)). In the present study, only a few lipid species showed significant differences between infected and mock-infected samples; this may be because *in vivo* bulk lipidomic analysis is not comparable to an *in vitro* approach. It has been suggested that the dysregulation of the polyunsaturated and saturated/monounsaturated lipid species could reflect the need for increased membrane fluidity for viral replication (114); however, the increase in PUFAs could also be a response to oxidative stress, as it is known that PUFAs can scavenge free radicals (115). The positive enrichment of GO terms associated with nitric oxide metabolic process/biosynthetic process, superoxide anion generation, superoxide metabolic process, respiratory burst, oxidoreductase activity, reactive oxygen species (ROS) metabolic process, (cellular) response to ROS, NADPH oxidase complex, (cellular) response to hydrogen peroxide, antioxidant activity, and cellular oxidant detoxification in the transcriptome/proteome provides evidence of an oxidative environment in the infected brains, possibly as a consequence of leukocyte activation (30). Additionally, infected brains showed a significantly increased abundance of quinic acid and urate, as already mentioned, two molecules with antioxidant properties (110, 116).

In the lipidome, functional enrichment analysis showed the positive enrichment of the cellular component lipid droplet (LION ontology (69)). Infection with SARS-CoV-2 and other +ssRNA viruses has been reported to increase lipid droplets in infected cells, as these support viral replication and are suggested to be involved in proinflammatory and antiviral responses (117–119). SARS-CoV-2 induces lipid droplet formation via activation and nuclear translocation of SREBP-1 and SREBP-2 (117). However, based on the absence of a significant difference in the abundance of triacylglycerols (TGs), one of the major component of lipid droplets (117), between the infected and mock-infected cohorts, and the absence of clearly identifiable lipid droplets at the ultrastructural level in infected neurons, we cannot be certain of their formation in infected brains.

Previous studies reported neuronal degeneration and necrosis/apoptosis in infected brains of K18-hACE2 mice intranasally challenged with ancestral SARS-CoV-2 strains at various doses (2.0x10^3^ to 1.0x10^6^ PFU; 1.0x10^3^ to 5.0x10^4^ TCID_50_), though generally higher than the Delta inoculation dose of 10^2^ PFU used in the present study (15, 17, 19, 23–28). The most convincing morphological changes at the light microscopy level in the present study were cytoplasmic vacuoles/clear spaces in infected neurons immunolabelled for viral NP, a feature that was also described in other studies (19, 23) and hypothesised to be related to viral replication (23). This aligns with our TEM results which suggest that these structures could correspond to distended ER/EGIC/lumen of the NE and CMs/LVCV. Our ultrastructural examination also detected the death of individual infected neurons. This was mainly indicated by nuclear pyknosis, with occasional evidence of phagocytosis of infected cells, but without satellitosis and overt neuronophagia as expected with neuronal necrosis due to viral infection. Our observation aligns with findings from another study which reported microglia in contact with and engulfing infected neurons (27). While our multi-omics approach did not allow us to determine the exact mechanisms leading to neuronal death, the transcriptome revealed positive enrichment of terms related to programmed necrotic cell death, pyroptotic cell death, apoptotic signalling pathway, and regulated necrosis. Several studies have shown that SARS-CoV-2 can interact with multiple cell death pathways, such as apoptosis, necroptosis, and pyroptosis (reviewed in (120, 121)); in K18-hACE2 mice, *in vitro* work on isolated neurons identified activation of the necroptosis pathway upon SARS-CoV-2 infection (13). Based on our ultrastructural findings, we suspect that neuronal death results from cellular exhaustion by the infection, rather than from direct cytopathic processes induced by the virus. This is further supported by the fact that the dead cells that can be readily identified at the light microscopic level in infected brains were occasional infiltrating leukocytes undergoing apoptosis (30); this was accompanied by the enrichment of leukocyte-related cell death terms (including lymphocyte (T/B) and neutrophilic leukocyte death) in the transcriptome (30).

Our ultrastructural investigations found evidence of viral particle release from infected neurons. Given the route of infection and the viral target cells in the brain, we expected to find evidence of neuron-to-neuron transmission. However, although we could demonstrate viral NP not only in neuronal cell body but also in cell processes, we did not find clear evidence of the latter and could not confirm the suspected mode of virus spread.

While neuronal infection did not obviously affect neuronal vitality, the extent of the viral replication-related ultrastructural changes in infected neurons suggests their exhaustion; we therefore expected at least evidence of impaired function. Indeed, the transcriptome showed negative enrichment of terms related to neurons, synapses, and behaviours. However, for several GABA and glutamate neurotransmitter receptors, the transcriptomics and proteomics datasets were discordant. Similarly, the metabolome showed no significant differences in GABA and glutamate between the two cohorts. Cell deconvolution suggested altered excitatory and inhibitory neuron proportions in infected brains, both lower than in the control group. Because the morphological results showed only negligible neuronal death, this difference is unlikely to reflect neuronal loss. However, SARS-CoV-2 suppresses host transcription and translation and induces host mRNA degradation; indeed, 60% of the RNA in an infected cell could be of viral origin (104, 105, 122). Therefore, we hypothesise that the reduced proportion of neurons in infected brains, as indicated by transcriptome-based cell deconvolution, reflects reduced transcript abundance.

In agreement with previous work by our group and others (14, 15, 19–21, 26–28, 123), we found morphological and molecular evidence of a glial response to neuronal infection, involving both the microglia and astrocytes. In regions of neuronal infection, most microglial cells exhibited a reactive “morphotype”, with increased soma area and thicker processes (45). These morphological changes are associated with altered microglial function, including surveillance for DAMPs and PAMPs, cytokine production, and phagocytosis of cellular debris (45), which aligns with the changes observed in our studies (30). Despite clear differences in microglial morphology between the Delta-infected and mock-infected cohorts, as expected (124), quantification of IBA1 tissue expression showed no significant differences between the two cohorts. Using *in situ* single-cell microglial analysis techniques, we found statistically significant differences in several morphological parameters between the Delta-infected and mock-infected cohorts, including a higher number of IBA1-positive cells in the infected brains, which aligns with the transcriptome-based cell deconvolution results. However, despite assessing 51 parameters for each IBA1-positive cell, the clustering approach could not recapitulate the observed change in microglial morphology. We suspect this is partly due to the inherent limitations of our automated image analysis pipeline, such as excluding processes of “normal” microglial cells, dataset bias towards the reactive “morphotype”, and overlap between microglial cells and monocytes/macrophages. Despite these limitations, our pipeline indirectly captured some of the differences observed at light microscopy. For example, in the infected brains, some detections were more likely to be just swollen cell processes rather than a microglial cell with a soma and cell processes. While this might have artificially increased the number of microglial cells in the infected brains, it aligns with our description that microglial cells in those brains have thicker cell processes.

Morphological changes in astrocytes in the infected brains were more subtle and could not be appreciated at the light microscopy level, including with IH for GFAP. This is not surprising since GFAP expression patterns do not correlate with the degree of astrocyte reactivity, nor does astrocyte morphology correlate with functional phenotypes (101). However, ultrastructurally, we detected astrocytes with distended cell bodies and processes, phagocytosis of cell debris, and increased glycogen, all features of reactive astrocytes (101). This aligns with our previous finding of an increase in glycogen in astrocytic endfeet in these brains (30); we consider this evidence of higher glucose demand by infected neurons.

This study has some potential limitations. We investigated animals at only one time point post-infection and challenged them with a single low dose. We chose this because we wanted to assess the brain and the effect on neurons in the worst-case scenario, and other studies have already investigated the longitudinal aspect of the infection (15, 17, 19), as well as a possible dose-effect (Supplemental Table S1). We also examined only female mice to reduce the number of confounding variables; as we were interested in the general mechanisms of neuronal infection and associated effects. Finally, we used a bulk-omic approach, which does not allow us to assign the observed changes to a specific cell type. However, since this was accompanied by in-depth *in situ* investigations, we were able to hypothesise which changes were likely related to neurons, and which ones to glial cells.

Taken together, our results confirm that in the K18-hACE2 model, intranasal challenge with the SARS-CoV-2 Delta variant infects only neurons, which support substantial viral replication that can lead to cell exhaustion and death. This suggests that in this murine model, SARS-CoV-2 behaves as a neurotropic, but not a classical neuropathic virus. Neuronal infection is associated with an innate immune and antiviral response and a limited reactive glial response. While the virus interferes with neuronal transcription and translation, the exact consequences on neuronal function, including beyond the acute phase, remain to be determined. Indeed, while some have claimed that K18-hACE2 mice infected with SARS-CoV-2 ancestral strains can survive the acute phase of infection (123, 125), none of these studies provided evidence that the surviving animals experienced brain infection at any point during the acute phase. Therefore, whether or not the outcome of brain infection is always lethal or remission can occur, remains unknown and requires further investigation. Nonetheless, the absence of overt neuronal cell death, despite widespread neuronal infection and viral replication, makes the intranasally challenged K18-hACE2 mouse model a candidate for investigating host-virus interactions, including the features that distinguish it from infections that cause overt neuronal death.

## Ethics approval and consent to participate

Animal work was approved by the local University of Liverpool Animal Welfare and Ethical Review Body and performed under UK Home Office Project Licence PP4715265. It was undertaken in accordance with locally approved risk assessments and standard operating procedures.

## Availability of data and materials

All data used for the generation of figures and the lists of genes of interest will be provided as supplemental data.

The code used for the -omic analyses will be uploaded to a public repository after publication.

Sequencing data has been deposited on Sequence Read Archive (SRA) (126) under the BioProject: PRJNA1220666. The mass spectrometry proteomics data have been deposited to the ProteomeXchange Consortium via the PRIDE (127) partner repository with the dataset identifier PXD074752. The mass spectrometry metabolomic and lipidomic data have been deposited to the MetaboLights repository (128) with the identifier REQ20260518219711.

The scripts used for the digital image analysis pipeline, as well as instructions on how to reproduce it, will be uploaded to a public repository after publication.

## Authors’ contribution

AK (Anja Kipar), JPS, UH: conception and design of the project. SDN, BH, RPR, AK (Adam Kirby), PS, LN, ME, MZ, AJ, FP, CRC: methodology. PS, AK (Adam Kirby), DM: *in vivo* experimental work. SDN, RPR, PS, BH, AK (Adam Kirby), LN, ME: data collection. SDN, RPR, LN, AK (Anja Kipar), UH, MZ, AJ, AO: data analysis. SDN, AK (Anja Kipar): writing of the original draft. All authors: review and editing. AK (Anja Kipar), JPS, EE: funding acquisition. AK (Anja Kipar): supervision of the project. All authors read and approved the manuscript.

## Acknowledgements

We are grateful to the technical staff in the Histology Laboratory, Institute of Veterinary Pathology, Vetsuisse Faculty, University of Zurich (IVPZ), for excellent technical support. We thank Dr Giuseppe Balistreri, Department of Virology, Faculty of Medicine, University of Helsinki, for fruitful discussions regarding the ultrastructural findings. We thank Dr Ludovico Silvestri and his team at Clepio Biotech Srl, Florence, Italy, for the acquisition of the 3D fluorescence imaging data. We would like to thank Professor Wei-Chung Cheng and Dr. Pei-Chun Shen at Program for Cancer Molecular Biology and Drug Discovery, College of Medicine, China Medical University, for their help in applying their R package, LipidSigR, to our analysis. We are grateful to Professor Sten Linnarsson and Peter Lönnerberg at the Laboratory of Molecular Neurobiology, department of Medical Biochemistry and Biophysics, at the Karolinska Institute, for their help in using their single cell transcriptomic data to build a single cell signature gene matrix for murine thalamus to be used for cell deconvolution on our bulk transcriptomic data. We would like to thank Professor Maria Lopez Ávalos, Department of Cellular biology, Genetic, and physiology, University of Malaga, for her assistance in the implementation of the FracLac analysis. We are grateful to Dr Pete Bankhead, Centre for Genomic & Experimental Medicine, Institute of Genetics and Cancer, The University of Edinburgh, for his assistance in the optimisation of the script of the IBA1 cells analysis in QuPath, and to Dr Flurin Sturzenegger, Center for Microscopy and Image Analysis, University of Zurich, for his support in the implementation of an automate pipeline for the Sholl analysis. We thank Dr Julien Boccard, School of Pharmaceutical Sciences, University of Geneva, and Dr Florence Mehl, Swiss Institute of Bioinformatics, for their help in the implementation and interpretation of the multiblock (low-level multi-omics) analysis.

