## Supplemental Material for "Widespread SARS-CoV-2 brain infection after intranasal challenge leads to innate and anti-viral immune response and interference with neuronal function in K18-hACE2 mice"

**Supplemental Tables**

**Supplemental Table S1**. Overview of the published studies investigating brain infection in K18-hACE2 mice, with information on route of infection, SARS-CoV-2 strain/VOC, inoculation dose, viral detection methods, and reported infected cell types in the brain. For listed references, see pages 28-36 of this document.

| SARS-CoV-2 strain/VOC | Dose | Route of infection | Mice | Virology | Infected cell type (brain) | Reference |
|---|---|---|---|---|---|---|
| 2019n-CoV/USA-WA1.2020 | $2.5x10^4$ PFU | IN | A | VL | Neuron[a, 1] | (1) |
| 2019n-CoV/USA-WA1.2020 | $1.0x10^4$ PFU<br>$1.0x10^6$ PFU | IN | A | VL, VT | Neuron[a,b] | (2) |
| human/ITA/Milan-UNIMI-1/2020 | $1.0x10^5$ $TCID_{50}$ | IN | B | VL, VT | Neuron[c,d]<br>Astrocyte[c,d]<br>Microglia[c,d] | (3) |
| human/ITA/Milan-UNIMI-1/2020 | $1.0x10^5$ $TCID_{50}$ | Aerosol | B | NBI | NBI | |
| hCoV-19/Singapore/2/2020 (SG12) | $2.0x10^4$ PFU<br>$2.0x10^5$ PFU | IN | C | VT | NA | (4) |
| USA-WA1/2020 | $1.0x10^5$ PFU | IN | B | VL, VT | NA | (5) |
| B.1 | $1.0x10^3$ $TCID_{50}$ | IN | B | VL | Neuron[d] | (6) |
| B.1.315/Beta | $1.0x10^3$ $TCID_{50}$ | IN | B | VL | Neuron[d] | |
| B.1.617.2/Delta | $1.0x10^3$ $TCID_{50}$ | IN | B | VL | Neuron[d] | |
| BA.1.1/Omicron | $1.0x10^3$ $TCID_{50}$ | IN | B | NBI | NBI | |
| hCoV-19/Spain/CT-2020030095/2020 | $1.0x10^4$ $TCID_{50}$ | IN | B | VL, VT | Neuron[d] | (7) |
| hCoV-19/Spain/CT-2020030095/2020 | $1.0x10^3$ $TCID_{50}$ | IN | B | VL, VT | Neuron[d] | |
| nCoV-WA1-2020 | $1.0x10^4$ $TCID_{50}$<br>$1.0x10^5$ $TCID_{50}$ | IN | B | VL, VT | Neuron[d] | (8) |
| USA-WA1/2020 | $1.0x10^4$ PFU | IN | B | VT | NA | (9) |
| NMDC10013001 | $1.0x10^2$ $TCID_{50}$ | IN | D | VL | Neuron[e]<br>Endothelial cells[e] | (10) |
| USA-WA1/2020 | $1.0x10^5$ PFU | IN | B | VT | Neuron[c]<br>Choroid plexus epithelium[c] | (11) |
| B.1.1.7 hCoV19/England/204820464/2020 (Alpha) | $1.0x10^3$ PFU | IN | B | VL | NA | (12) |
| B1.617.2 hCoV-19/USA/WV-WVU-WV118685/2021 (Delta) | $1.0x10^3$ PFU | IN | B | VL | NA | |
| Hu-1 (Wuhan, early strain) | $2.5x10^4$ $TCID_{50}$ | IN | B | VL, VT | Neuron[c,d] | (13) |
| B1.617.2 (Delta) | $2.5x10^4$ $TCID_{50}$ | IN | B | VL, VT | Neuron[c,d] | |
| hCoV-2/human/Liverpool/REMRQ0001/2020 | $1.0x10^3$ PFU | IN | B | VL | Neuron[d] | (14) |
| hCoV-2/human/Liverpool/REMRQ0001/2020 | $1.0x10^4$ PFU | IN | B | VL | Neuron[d] | |
| B.1.617.2 hCoV/England/SHEF-10E8F3B/2021 (Delta) | $1.0x10^2$ PFU | IN | B | VL | Neuron[d] | |

| | | | | | | |
|---|---|---|---|---|---|---|
| B.1.617.2 hCoV/England/SHEF-10E8F3B/2021 (Delta) | $1.0x10^3$ PFU | IN | B | VL | Neuron[d] | |
| hCoV-2/human/Liverpool/REMRQ0001/2020 | $1.0x10^3$ PFU | IN | B | VL | Neuron[a,d] | (15) |
| hCoV-2/human/Liverpool/REMRQ0001/2020 | $1.0x10^4$ PFU | IN | B | VL | Neuron[a,d] | |
| B.1.1.7 Human nCoV19 isolate/England/202012/01B (Alpha) | $1.0x10^3$ PFU | IN | B | VL | Neuron[a,d] | |
| B.1.351 20I/501.V2.HV001 (Beta) | $1.0x10^3$ PFU | IN | B | VL | Neuron[a,d] | |
| B.1.617.2 hCoV/England/SHEF-10E8F3B/2021 (Delta) | $1.0x10^3$ PFU | IN | B | VL | Neuron[a,d] | |
| B.1.1.529/BA.1 M21021166 (Omicron) | $1.0x10^3$ PFU | IN | B | NBI | NBI | |
| NCCP 43326, S type | $1.0x10^6$ PFU | IN | B | NA | Neuron[d] | (16) |
| NCCP 43326, S type | $1.0x10^6$ PFU | Inhalation | B | NA | Neuron[d] | |
| human/USA/WA-CDC-WA1/2020 | $2.0x10^1$ PFU<br>$2.0x10^2$ PFU<br>$2.0x10^3$ PFU<br>$2.0x10^4$ PFU | IN | B | VL | Neuron[d] | (17) |
| 2019n-CoV/USA-WA1/2019 | $1.0x10^5$ PFU | IN | B | VT | Neuron[d, 1] | (18) |
| BavPat1/2020 | $5.0x10^4$ $TCID_{50}$ | IN | B | VL | Neuron[d] | (19) |
| BetaCoV/Korea/KCD03/2020 (wild-type) | $2.0x10^3$ PFU | IN | B | NA | Neuron[c]<br>Astrocyte[c]<br>Oligodendrocyte[c] | (20) |
| BetaCoV/Korea/KCD03/2020 (wild-type) | $2.0x10^3$ PFU | IN | B | VL | NA | (21) |
| B.1.351 hCoV-19/South Korea/KDCA0463/2020 (Beta) | $2.0x10^3$ PFU<br>($1.0x10^4$ PFU) | IN | B | VL | NA | |
| NCCP 43326, S type | $1.0x10^5$ PFU | IN | B | NA | Neuron[a,c] | (22) |
| hu/USA/CA-CZB-59X002/2020 (MT394529) B.1 (WA1-like) | $1.0x10^4$ PFU | IN | B | VT | Neuron[c] | (23) |
| hu/USA/CA_CDC_5574/2020 B.1.1.7 (Alpha) | $1.0x10^4$ PFU | IN | B | VT | Neuron[c] | |
| hCoV-19/USA/MD-HP01542/2021 B.1.351 (Beta) | $1.0x10^4$ PFU | IN | B | VT | Neuron[c] | |
| hCoV-19/USA/PHC658/2021 B.1.617.2 (Delta) | $1.0x10^4$ PFU | IN | B | VT | Neuron[c] | |
| hCoV-19/USA/HI-CDC-4359259-001/2021 B.1.1.529 (Omicron) | $1.0x10^4$ PFU | IN | B | NBI | NBI | |
| hCoV-19/USA/MD-HP40900/2022 B.1.1.529 (Omicron XBB.1.5) | $1.0x10^4$ PFU | IN | B | VT | Neuron[c] | |
| USA-WA1/2020 | $1.5x10^6$ PFU | IN | B | VL, VT | Neuron[f] | (24) |
| WA-1/2020 | $2.0x10^3$ PFU<br>$2.0x10^4$ PFU | IN | B | NA | Neuron[g,h] | (25) |
| hCoV-2/human/Liverpool/REMRQ0001/2020 | $1.0x10^4$ PFU | IN | B | NA | Neuron[d] | (26) |
| hCoV-2/human/Liverpool/REMRQ0001/2020 | $1.0x10^4$ PFU | IN | E | VL | NA | (27) |
| hCoV-2/human/Liverpool/REMRQ0001/2020 | $1.0x10^3$ PFU | IN | E | VL | NA | |

| | | | | | | |
|---|---|---|---|---|---|---|
| B.1 (Wuhan) | $1.0x10^4$ PFU | IN | A | VL | NA | (28) |
| B.1.1.7 (Alpha) | $1.0x10^4$ PFU | IN | A | VL | NA | |
| B.1.351 (Beta) | $1.0x10^4$ PFU | IN | A | VL | NA | |
| B.1.617.2 (Delta) | $1.0x10^4$ PFU | IN | A | VL | NA | |
| B.1.529 (Omicron) | $1.0x10^4$ PFU | IN | A | VL | NA | |
| human/KOR/KCDC03/2020 | $1.0x10^4$ PFU | IN | B | VL | NA | (29) |
| human/KOR/KCDC03/2020 | $1.0x10^4$ PFU | IT | B | VL | NA | |
| human/KOR/KCDC03/2020 | $1.0x10^4$ PFU | IC | B | VL | NA | |
| human/KOR/KCDC03/2020 | $1.0x10^4$ PFU | Eye-drop | B | NBI | NA | |
| human/KOR/KCDC03/2020 | $1.0x10^4$ PFU | IV | B | VL | NA | |
| USA-WA1/2020 | $1.0x10^5$ PFU | IN | B | VL | NA | (30) |
| USA-WA1/2020 | $1.0x10^4$ PFU - $1.0x10^5$ PFU | IN | B | VL | Neuron[a] | (31) |
| USA-WA1/2020 | $1.0x10^4$ $TCID_{50}$ | IN | B | VL | NA | (32) |
| New York-PV09158/2020 lineage B.1.3 | $1.0x10^3$ $TCID_{50}$ | IN | B | VL | NA | |
| USA/CA_CDC_5574/2020 lineage B.1.1.7 (Alpha) | $1.0x10^2$ $TCID_{50}$ | IN | B | VL | NA | |
| hCoV-19/South Africa/KRISP-EC-K005321/2020 lineage B.1.351 (Beta) | $1.0x10^2$ $TCID_{50}$ | IN | B | VL | NA | |
| WH-09/human/2020/CHN | $1.0x10^5$ $TCID_{50}$ | IN | D | VL | Neuron[b, 1] | (33) |
| USA-WA1/2020 (wild type) | $1.0x10^4$ PFU | IN | B | VL | Neuron[d, 1] | (34) |
| B.1.617.2 (Delta) | $1.0x10^4$ PFU | IN | B | VL | Neuron[d, 1] | |
| B.1.1.529 (Omicron) | $1.0x10^4$ PFU | IN | B | VL | NA | |
| human/KOR/KCDC03-NCCP43326/2020 | $2.0x10^4$ PFU | IN | B | VL | Microglia[c,h] | (35) |
| MAD6 | $1.0x10^5$ PFU | IN | B | VL | Neuron[c,d]<br>Endothelial cells[c,d]<br>Microglia[c,d] | (36) |

**Abbreviations:** IN – intranasal; IC – intracerebral; IT – intratracheal; IV - intravenous NA: not available; NBI: no brain infection; PFU: plaque-forming unit; $TCID_{50}$: median tissue culture infectious dose; VL: viral loads; VT: viral titers.

**Mice**: A - 2B6.Cg-Tg(K18-ACE2)2Prlmn/J; B - B6.Cg-Tg(K18-ACE2)2Prlmn/J; C - B6;SJL-Tg(K18-hACE2)2Prlmn/J; D - C57BL/6JGpt-H11em1Cin(K18-hACE2)/Gpt; E - Heterozygote hACE2-transgenic C57BL/6

**Method to detect infection**: a – RNA-*in situ* hydridization (ISH) for SARS-CoV-2 spike protein (S); b - immunohistochemistry (IH) for SARS-CoV-2 S; c - immunofluorescence for SARS-CoV-2 nucleocapsid protein (NP); d - IH for SARS-CoV-2 NP; e – Fluorescence ISH for SARS-CoV-2 S; f - iDISCO+ whole brain immunolabeling (light sheet microscopy) for SARS-CoV-2 NP; g – RNA-ISH for SARS-CoV-2 genomic RNA; h - IF for SARS-CoV-2 S; 1: Neurons were identified as the infected cell type based on the figures, although the authors did not state in the text which cells were infected.

**Supplemental Table S2**. Study cohorts and investigations undertaken.

| Cohort | Virus | PFU/mouse | DPI: animal no | Cohort in (14) | Investigations |
|---|---|---|---|---|---|
| 1 | Delta | $10^2$ | **6**: 1.1 - 1.3<br>**7**: 1.4 - 1.8 | 2.1.1 – 2.1.3<br>2.1.4 – 2.1.8 | H; IH; RT-qPCR; multi-omics*; TEM, immune-EM |
| 2 | Mock | - | **7**: 2.1 - 2.6 | 2.3.1 – 2.3.6 | H; IH**; multi-omics; TEM, immune-EM |

**Abbreviations**: Animal no: animal number for identification of individual mice; DPI: days post infection (day of euthanasia); Delta: SARS-CoV-2 Delta isolate; H: histological examination of the brain (HE-stained section); IH: immunohistochemistry for SARS-CoV-2 nucleocapsid protein (NP), IBA1 and GFAP; immune-EM: immune electron microscopy for SARS-CoV-2 N protein performed on the brain; PFU: plaque-forming units; Mock: mock-infected; multi-omics: bulk transcriptomic, proteomic, metabolomic, and lipidomic performed on the brain; RT-qPCR: reverse transcriptase-quantitative PCR for SARS-CoV-2 N1 (26) performed on lung and brain tissue; TEM: transmission electron microscopy performed on the brain. *No transcriptomic analysis was performed for mouse 1.8. **Mice 2.2-2.5 brains were not available for IH for GFAP and IBA1.

**Supplemental Table S3.** Suggested neuron "phenotypes" based on stages of SARS-CoV-2 replication in neurons, as indicated by the presence of ultrastructural features and immune-EM for viral NP; with reference to the figures illustrating the relevant features.

| Stage of viral replication | Ultrastructural features in infected neurons | Figure |
|---|---|---|
| 1. Early replication | - Occasional DMVs | 2A,<br>S2B |
| 2. Advanced replication | - Increased numbers of DMVs<br>- Membrane-bound viral particles<br>- Aggregates of NP in the cytoplasm | 2B,<br>S2C,<br>S3A, |
| 3. Late replication | - Numerous DMVs<br>- Abundant membrane-bound viral particles<br>- Large aggregates of NP in the cytoplasm<br>- Single-membrane vacuoles of variable size, some containing viral particles and multilamellar structures<br>- Irregular nuclear contour | 1A,<br>2C,<br>S2D |
| 4. Exhausting viral replication | - Pronounced vacuolation of the cytoplasm<br>- DMVs, multilamellar structures, and viral particles within vacuoles<br>- Irregular nuclear contour | 1B-D,<br>2D |

**Abbreviations**: DMVs: double-membrane vesicles; NP: SARS-CoV-2 nucleocapsid protein.

**Supplemental Table S4**. Statistical analysis between the Delta-infected and mock-infected groups, all brain regions together (separate file). For each measurement of the image analysis pipeline, the p-value and details on the statistical test are shown.

**Supplemental Table S5**. Statistical analysis between the Delta-infected and mock-infected groups, Cerebral Nuclei/Interbrain (separate file). For each measurement of the image analysis pipeline, the per group median, p-value, and adjusted p-value are shown.

**Supplemental Table S6**. Statistical analysis between the Delta-infected and mock-infected groups, Midbrain/Hindbrain (separate file). For each measurement of the image analysis pipeline, the p-value and details on the statistical test are shown.

**Supplemental Table S7**. The median of each measurement of the image analysis pipeline, per cluster (separate file).

**Supplemental Table S8.** List of R packages used for the image analysis pipeline and the -omic analyses. For listed references, see pages 28-36 of this document.

A) Image analysis pipeline

| **Package** | **Version** | **Reference** |
|---|---|---|
| cluster | 2.1.8.1 | (37) |
| dplyr | 1.1.4 | (38) |
| factoextra | 1.0.7 | (39) |
| forcats | 1.0.1 | (40) |
| ggcorrplot | 0.1.4.1 | (41) |
| ggplot2 | 4.0.1 | (42) |
| ggrepel | 0.9.6 | (43) |
| patchwork | 1.3.2 | (44) |
| PCAtools | 2.22.1 | (45) |
| reshape2 | 1.4.5 | (46) |
| ropls | 1.42.0 | (47) |
| rstatix | 0.7.3 | (48) |
| Rtsne | 0.17 | (49) |
| stringr | 1.6.0 | (50) |
| tidyr | 1.3.2 | (51) |
| tidyselect | 1.2.1 | (52) |
| umap | 0.2.10.0 | (53) |

B) Omic analyses

| Package | Version | References |
|---|---|---|
| AnnotationDbi | 1.72.0 | (54) |
| clusterProfiler | 4.18.4 | (55–58) |
| ComplexHeatmap | 2.26.0 | (59) |
| ConsensusOPLS | 1.1.0 | (60) |
| cowplot | 1.2.0 | (61) |
| DEP2 | 0.5.28.2 | (62) |
| dplyr | 1.1.4 | (38) |
| edgeR | 4.8.2 | (63) |
| EnhancedVolcano | 1.28.2 | (64) |
| enrichplot | 1.30.4 | (65) |
| FELLA | 1.30.0 | (66) |
| fgsea | 1.36.2 | (67) |
| forcats | 1.0.1 | (40) |
| ggplot2 | 4.0.1 | (42) |
| ggridges | 0.5.7 | (68) |
| ggsci | 4.2.0 | (69) |
| GO.db | 3.22.0 | (70) |
| limma | 3.66.0 | (71) |
| LipidsigR | 1.0.4 | (72–74) |
| MBAnalysis | 2.2.0 | (75) |
| org.Mm.eg.db | 3.22.0 | (76) |
| patchwork | 1.3.2 | (44) |
| PCAtools | 2.22.1 | (45) |
| pheatmap | 1.0.13 | (77) |
| reactome.db | 1.95.0 | (78) |
| ReactomePA | 1.54.0 | (79) |
| RefMet | 1.0.0 | (80) |
| reshape2 | 1.4.5 | (46) |
| ropls | 1.42.0 | (47) |
| rstatix | 0.7.3 | (48) |
| stringr | 1.6.0 | (50) |
| tibble | 3.3.1 | (81) |
| tidyr | 1.3.2 | (51) |
| tidyselect | 1.2.1 | (52) |
| tximport | 1.32.0 | (82) |
| WGCNA | 1.74 | (83) |

**Supplemental Table S9.** Details of the clustering of the different cell types for the cell deconvolution using CIBERSORTx (separate file).

### Supplemental Figures

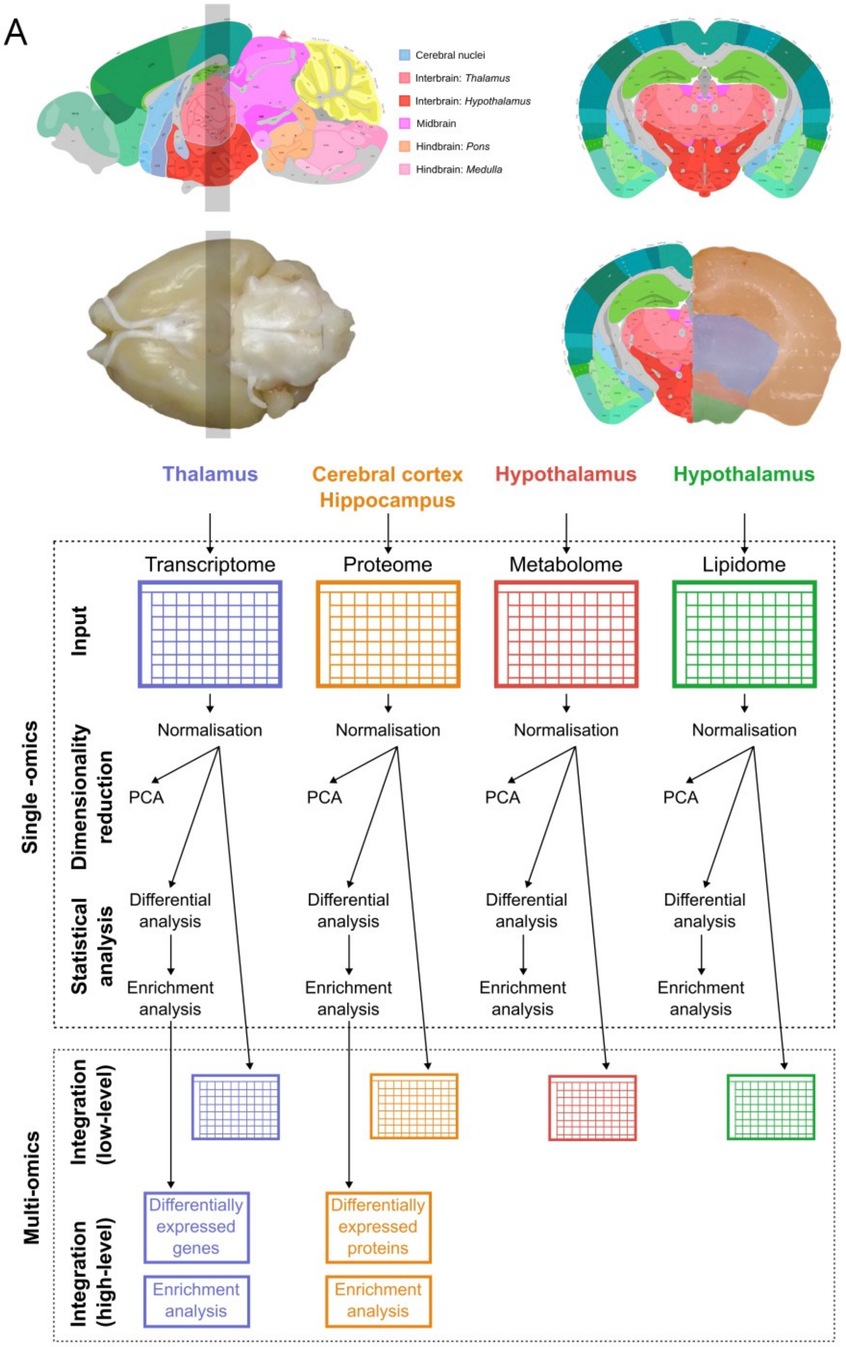


**Supplemental Figure S1.** Overview of the multi-omics pipeline and the image analysis workflow. **A)** Illustration of the tissue slice prepared by coronal sections (grey rectangle) to provide the samples for transmission electron microscopy and for the bulk transcriptomic, proteomic, metabolomic, and lipidomic analyses. Indicated are the annotated regions for quantifying SARS-CoV-2 NP, IBA1, and GFAP expression, namely: cerebral nuclei and interbrain; midbrain and hindbrain. Pictures of an example brain are mapped to the Allen Mouse Brain Atlas (Allen Reference Atlas – Adult Mouse Coronal Sections (image 75), Mouse, P56 Sagittal (image 21) - https://atlas.brain-map.org/) (84–87). **B)** Overview of the workflow used for the single -omics and the multi-omics analyses. For the lipidome, the positive and negative datasets were analysed separately. PCA: principal component analysis. Information on animal cohorts and analyses undertaken is provided in Supplemental Table S2. For listed references, see pages 28-36 of this document.

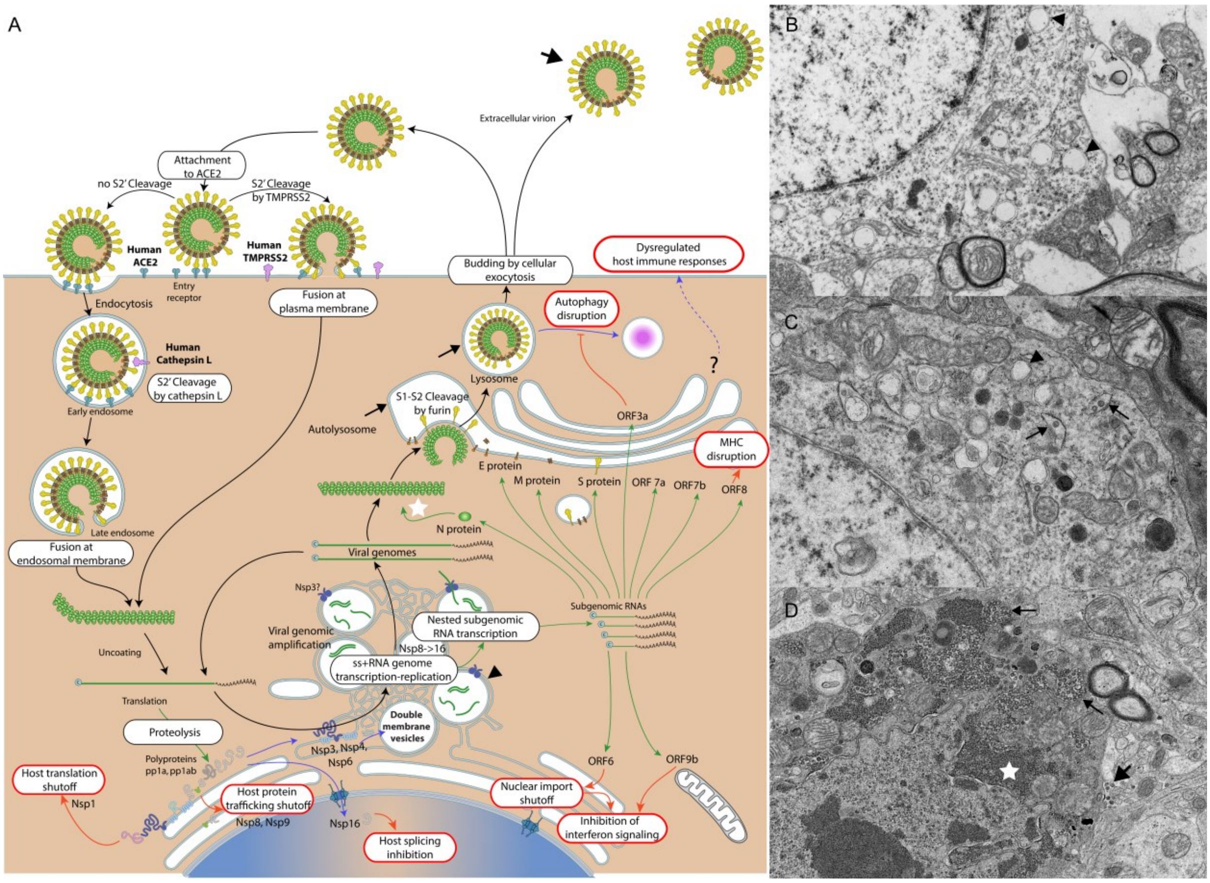


**Supplemental Figure S2. Brainstem, K18-hACE2 mice after intranasal challenge with SARS-CoV-2 Delta variant ($10^2$ PFU/mouse); 6-7 dpi.** Correlation between the SARS-CoV-2 replication cycle and key elements of the SARS-CoV-2 replication cycle observed at the ultrastructural level in infected neurons. **A)** Illustration of the coronavirus life-cycle (adapted from ViralZone (88) - https://viralzone.expasy.org/9096). **B)** Early viral replication, represented by occasional double-membrane vesicles (DMVs) (arrowhead) in the cytoplasm. **C)** Advanced viral replication, represented by DMVs (arrowhead) and membrane-bound viral particles (arrows) in the cytoplasm. **D)** Late viral replication, represented by abundant membrane-bound viral particles (arrows) and large aggregates of nucleocapsid protein (white star) in the cytoplasm. Viral particles in the extracellular space are visible (large arrow). Images B-D are identical to Figure 2A-C (higher magnifications). For listed references, see pages 28-36 of this document.

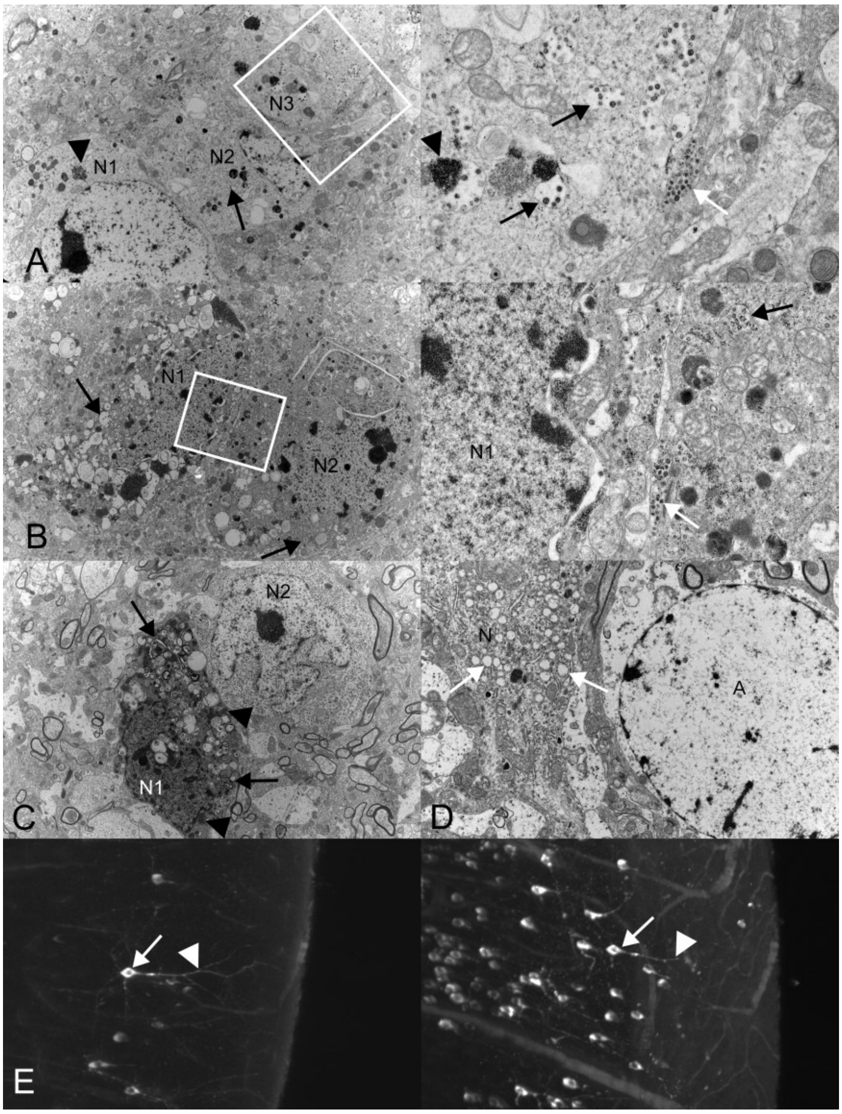


**Supplemental Figure S3. A-D) Brainstem, K18-hACE2 mice after intranasal challenge with SARS-CoV-2 Delta variant ($10^2$ PFU/mouse); 6-7 dpi. A, B)** Immune electron microscopy for SARS-CoV-2 nucleocapsid protein (NP). SARS-CoV-2 infection of neurons is proven by labelling of viral structures with an antibody directed against SARS-CoV-2 NP. The images on the right show closer views of the region indicated by the white rectangles in the images on the left. **A)** Three adjacent neurons (N1-3). Aggregates of NP (arrowheads, N1 and N2) and membrane-bound viral particles (black arrows) are present in the cytoplasm, consistent with advanced viral replication. The white arrow highlights viral particles in the extracellular space. **B)** Two adjacent neurons (N1, N2), both with membrane-bound viral particles (black arrows) in the cytoplasm. The white arrow highlights viral particles in the extracellular space between the infected neurons. **C, D)** Electron microscopy. **C)** Two neurons in direct contact, the left (N1) showing features of viral replication, with aggregates of NP (arrowheads) and accumulation of membranes (arrows), while the right (N2) is not infected. **D)** Uninfected astrocyte (A) immediately adjacent to an infected neuron (N) showing double-membrane vesicles (DMVs) in the cytoplasm (white arrow). **E) Cerebral cortex, K18-hACE2 mouse after intranasal challenge with SARS-CoV-2 Delta variant ($10^3$ PFU/mouse); 8 dpi.** Volumetric images of SARS-CoV-2 NP expression in a slice of cerebral cortex. The images on the left (attenuated maximum intensity projection (MIP); nearer positive neurons appear brighter) and the right (MIP; near and far positive neurons are rendered in a similar way) illustrate NP in neuronal bodies (arrow) and processes (arrowhead). The same neuron is indicated in both images.

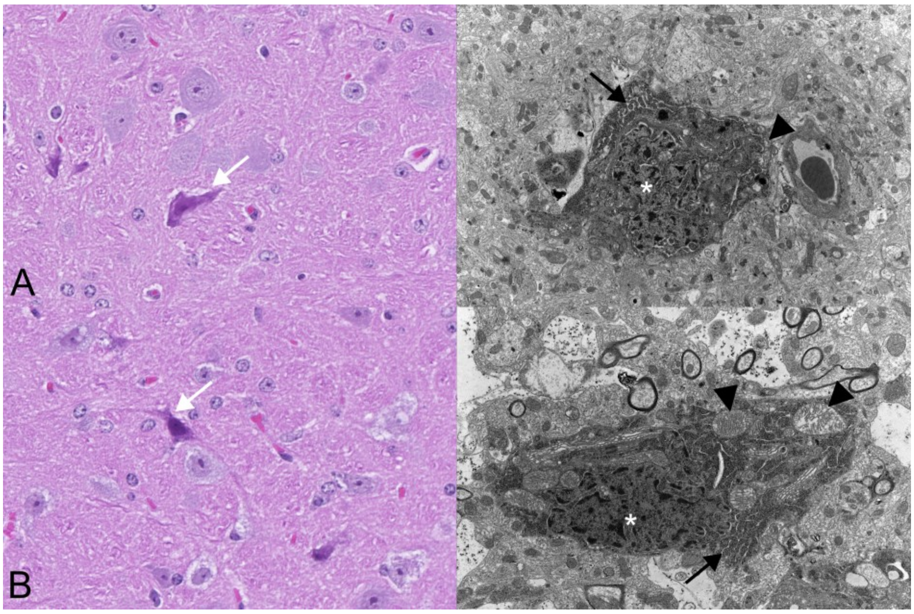


**Supplemental Figure S4. Brainstem, K18-hACE2 mice. A)** Mock-infected mouse. **B)** SARS-CoV-2 Delta variant ($10^2$ PFU/mouse) infected mice with widespread neuronal infection; 6/7 dpi. Left images: haematoxylin and eosin (HE) stained sections with individual dark neurons (white arrows), otherwise unaltered. Right: Transmission electron microscopy images of dark neurons (uninfected), with electron-dense cytoplasm containing distended endoplasmic reticulum (black arrows) and swollen mitochondria (arrowheads), and an electron-dense, shrunken nucleus with irregular contours (asterisks).

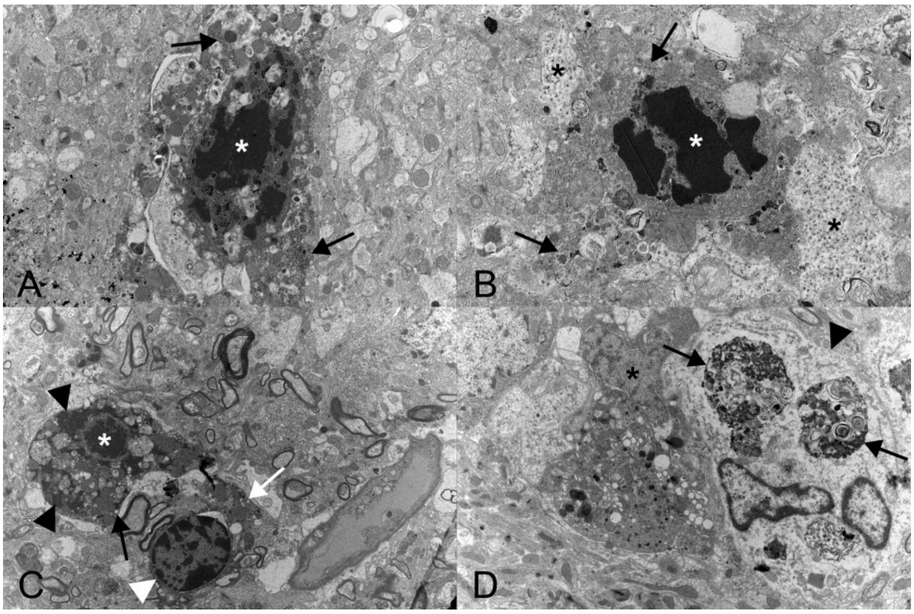


**Supplemental Figure S5. Brainstem, K18-hACE2 mice after intranasal challenge with SARS-CoV-2 Delta variant ($10^2$ PFU/mouse); 6-7 dpi. Ultrastructural evidence of cell death in infected neurons.** Nuclei exhibit angular electron-dense clumps of chromatin (white asterisks), consistent with pyknosis. **A)** Immune electron microscopy for SARS-CoV-2 nucleocapsid protein (NP) highlights membrane-bound viral particles in the cytoplasm (arrows). **B)** The cytoplasm contains membrane-bound viral particles (arrows). Adjacent to the infected neuron are swollen astrocytic processes (black asterisks). **C)** The cytoplasm contains membrane-bound viral particles (arrow) and NP aggregates (black arrowheads). Adjacent is a glial cell (white arrowhead) that contains phagocytosed cell debris (white arrow). **D)** Phagocytosed cell debris (arrows), including structures reminiscent of viral replication, are observed in the cytoplasm of a cell (arrowhead) with morphology consistent with an astrocyte. Immediately adjacent is an infected neuron (asterisk).

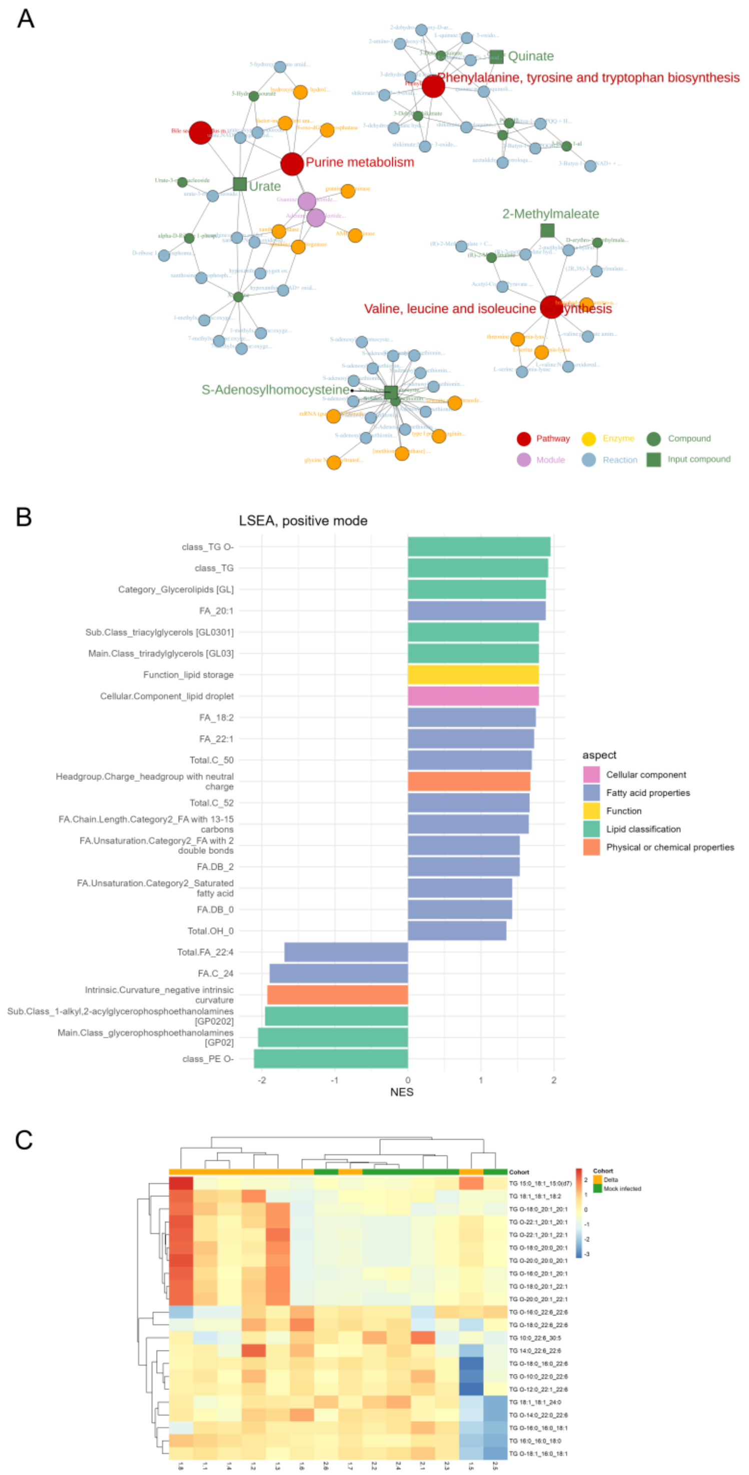


**Supplemental Figure S6. Metabolome and lipidome (hypothalamus), K18-hACE2 mice after intranasal challenge with SARS-CoV-2 Delta variant ($10^2$ PFU/mouse) and mock-infected; 6-7 dpi. Functional enrichment analysis**. **A)** Illustration of the significantly different metabolites mapped to a KEGG compound identifier, using FELLA. **B)** Illustration of the functional enrichment analysis for the lipidome (positive mode), using LSEA. NES: normalised enrichment score. **C)** Illustration of the abundance for the TG and TG O- lipid species.

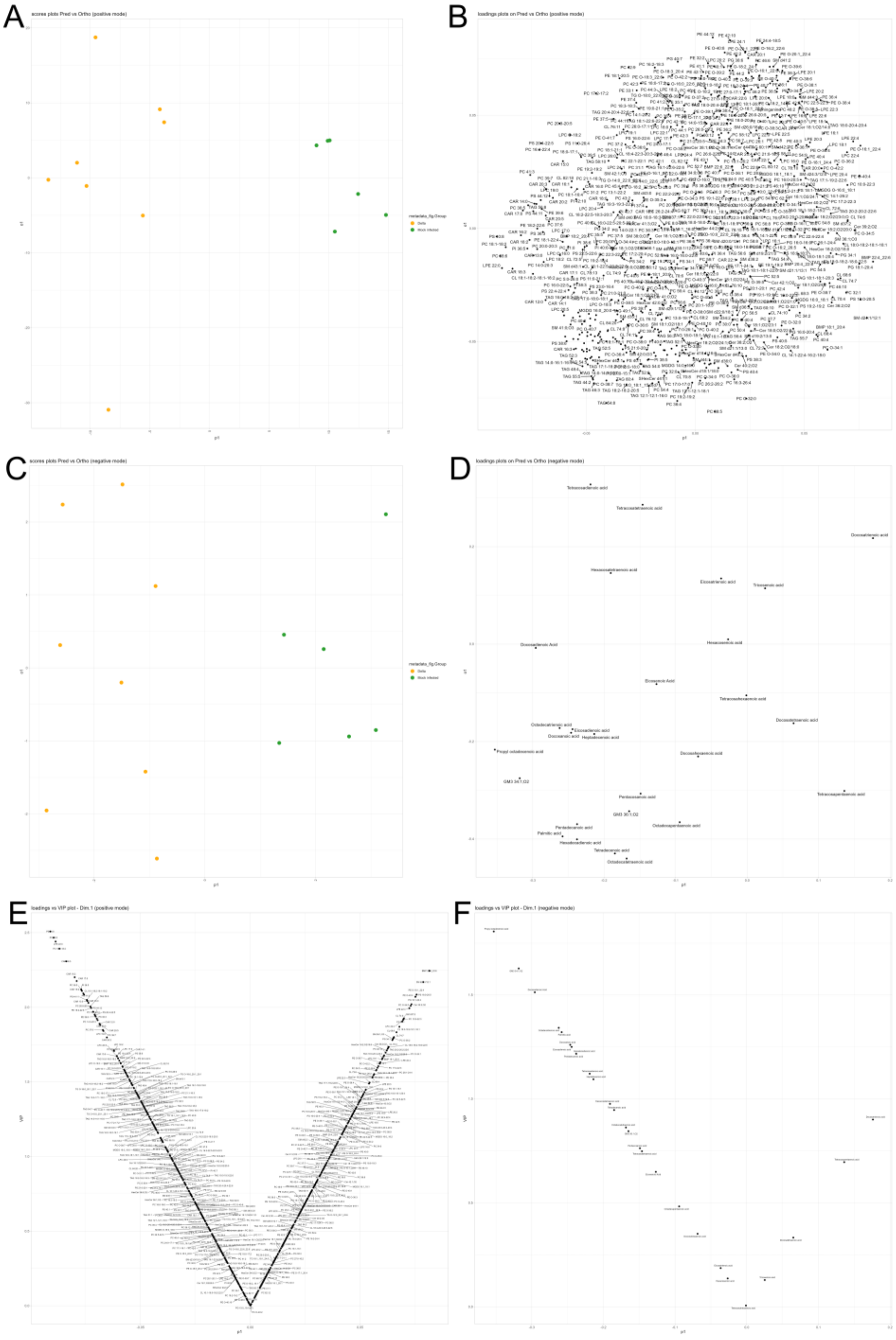


**Supplemental Figure S7. Lipidome (hypothalamus), K18-hACE2 mice after intranasal challenge with SARS-CoV-2 Delta variant ($10^2$ PFU/mouse) and mock-infected; 6-7 dpi. Orthogonal Partial Least Squares Discriminant Analysis (O-PLS-DA) showing the clustering of the samples based on a cohort discrimination. A, B)** Positive mode: illustration of the sample (A) and loading (B) plots. **C, D)** Negative mode: illustration of the sample (A) and loading (B) plots. **E, F)** Plot of the Variable Importance in Projection (VIP) for the positive (E) and negative (F) mode. The quality metrics of the models are available in Supplemental Material and Methods.

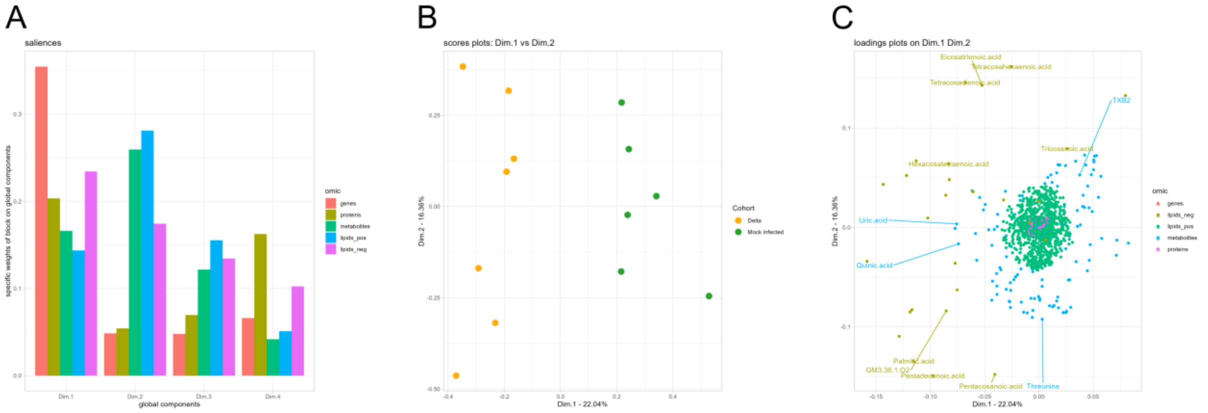


**Supplemental Figure S8. Multi-omics integration, low-level, Common Dimension method (ComDim). A)** Illustration of the contribution of each block (i.e. transcriptome (genes), proteome (proteins), metabolome (metabolites), lipidome (positive mode) (lipids_pos), lipidome negative mode (lipids_neg)) to the first four components. **B)** Illustration of the clustering of the samples into their respective cohort, namely SARS-CoV-2 Delta-infected (Delta) and mock-infected, along component 1 (Dim. 1) and component 2 (Dim. 2). **C)** Illustration of the loadings and their contribution to the clustering of the samples along components 1 and 2.

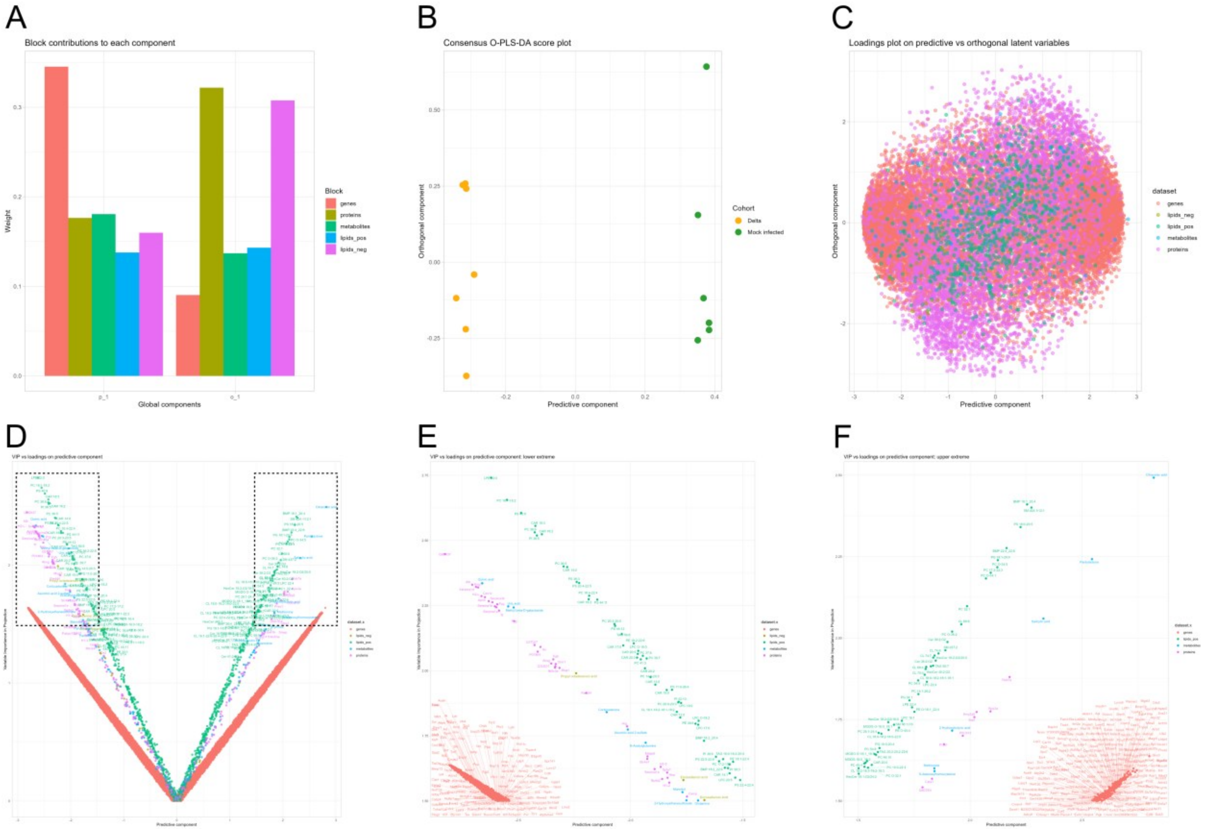


**Supplemental Figure S9. Multi-omics integration, low-level, consensus Orthogonal Partial Least Squares Discriminant Analysis (O-PLS-DA). A)** Illustration of the contribution of each block (i.e. transcriptome (genes), proteome (proteins), metabolome (metabolites), lipidome (positive mode) (lipids_pos), lipidome negative mode (lipids_neg)) to the global components. **B)** Illustration of the clustering of the samples into their respective cohort, namely SARS-CoV-2 Delta-infected (Delta) and mock-infected, along the predictive component and the orthogonal component. **C)** Illustration of the loadings and their contribution to the clustering of the samples along the predictive component and the orthogonal component. **D)** Illustration of the Variable Importance in Projection relative to the Predictive component. The variables at the two extremes of the plot, indicated by the dotted line boxes, are shown in E and F. **E)** Illustration of the Variable Importance in Projection relative to the Predictive component (left box in D). **F)** Illustration of the Variable Importance in Projection relative to the Predictive component (right box in D).

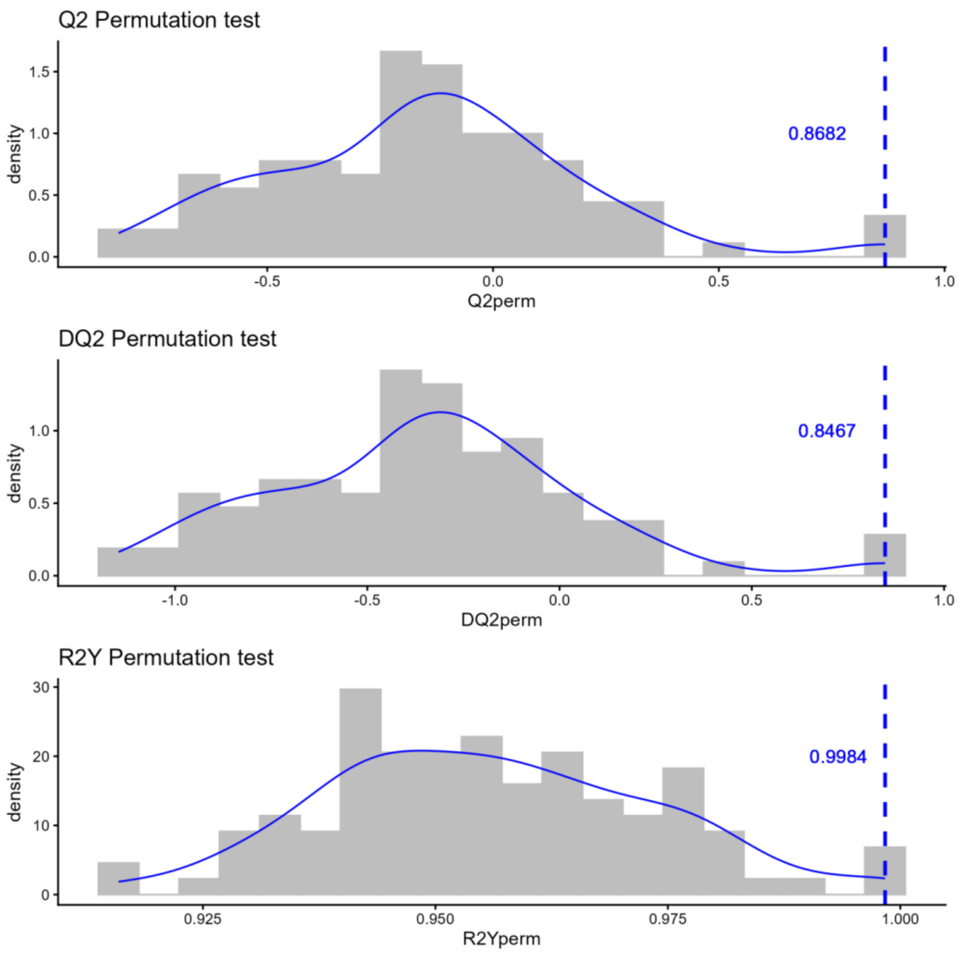


**Supplemental Figure S10**. **Multi-omics integration, low-level, consensus Orthogonal Partial Least Squares Discriminant Analysis (O-PLS-DA).** Distribution of permuted values (random permutations, n = 100, Leave-One-Out cross-validation) for: R-squared ($R^2Y$), Stone Geisser $Q^2$ (Q2), and discriminant $Q^2$ (DQ2) index.

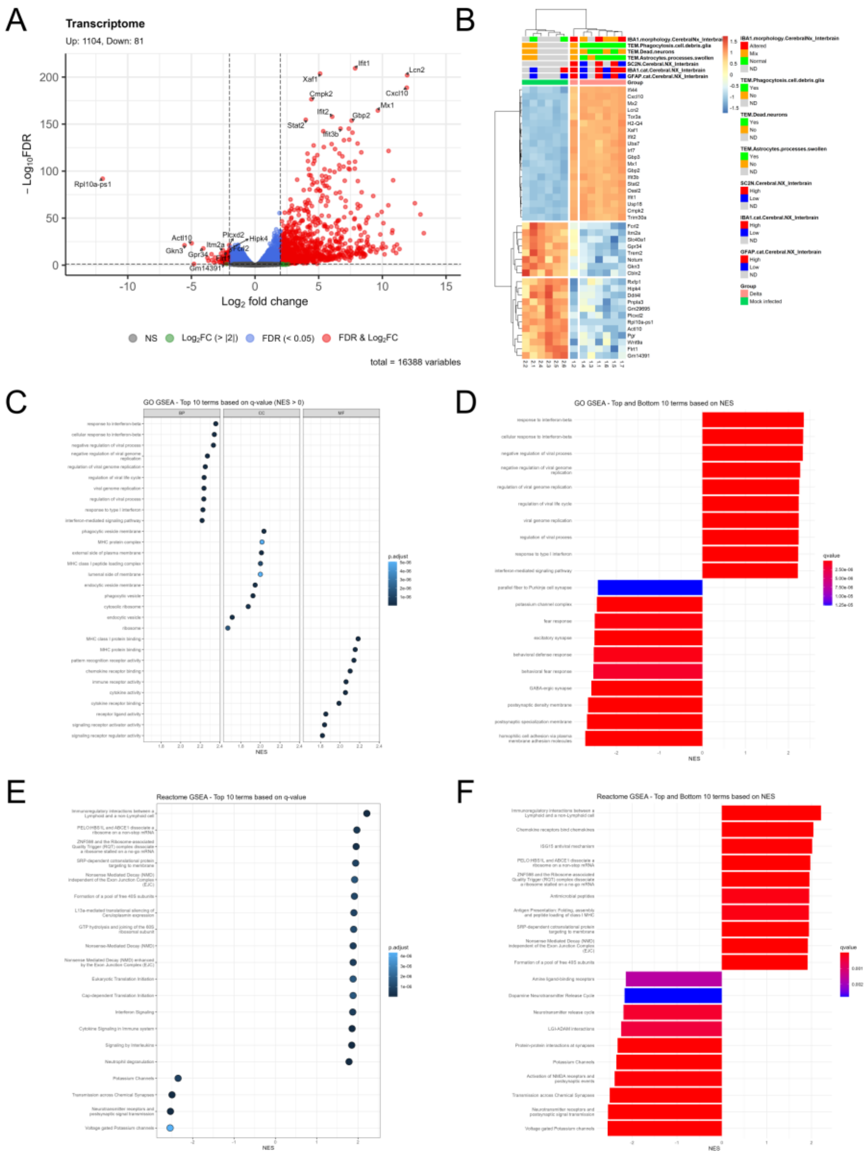


**Supplemental Figure S11. Transcriptome (thalamus), K18-hACE2 mice after intranasal challenge with SARS-CoV-2 Delta variant ($10^2$ PFU/mouse) and mock-infected; 6-7 dpi. A)** The Delta-infected (Delta) cohort shows significant up-regulation of genes; several of these are involved in inflammatory, innate immune, and antiviral responses. **B)** Heatmap showing the top and bottom (based on $log_2FC$) most significantly (FDR) changed genes. **C)** Top 10 positively enriched Gene Ontology (GO) terms, based on the q-value, for biological process (BP), cellular component (CC), and molecular function (MF). **D)** Top and bottom 10 most enriched GO terms, based on the normalised enrichment score (NES). **E)** Top 10 positively enriched Reactome terms, based on the q-value. **F)** Top and bottom 10 most enriched GO terms, based on NES. FDR: false discovery rate; $log_2FC$: $log_2$ fold change.

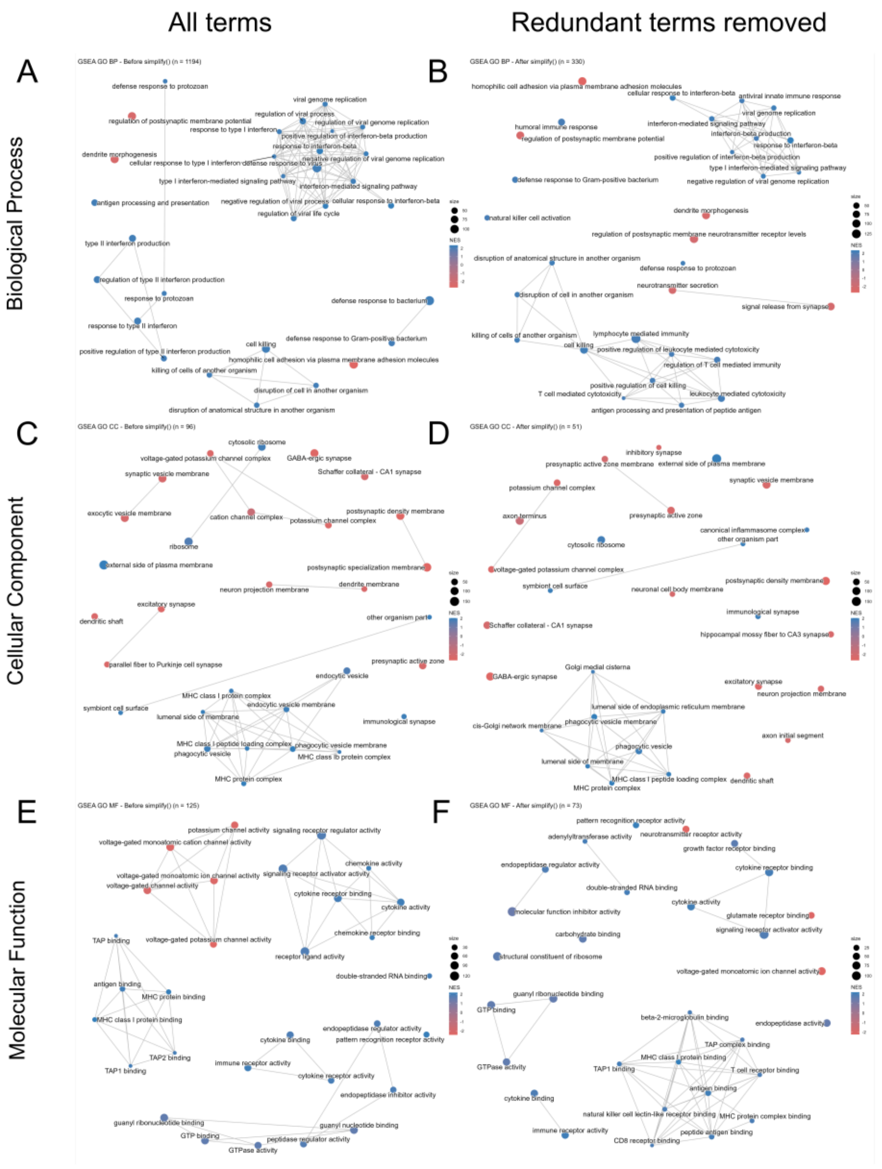


**Supplemental Figure S12. Transcriptome (thalamus), K18-hACE2 mice after intranasal challenge with SARS-CoV-2 Delta variant ($10^2$ PFU/mouse) and mock-infected; 6-7 dpi. Gene Ontology (GO) functional enrichment analysis.** Enrichment maps showing the top 30 enriched terms for the biological process (BP) (**A, B**), cellular component (CC) (**C, D**), and molecular function (MF) (**E, F**). In B, D, and F, we summarised the results of the enrichment analysis (A, C, E) using the clusterProfiler::simplify() function.

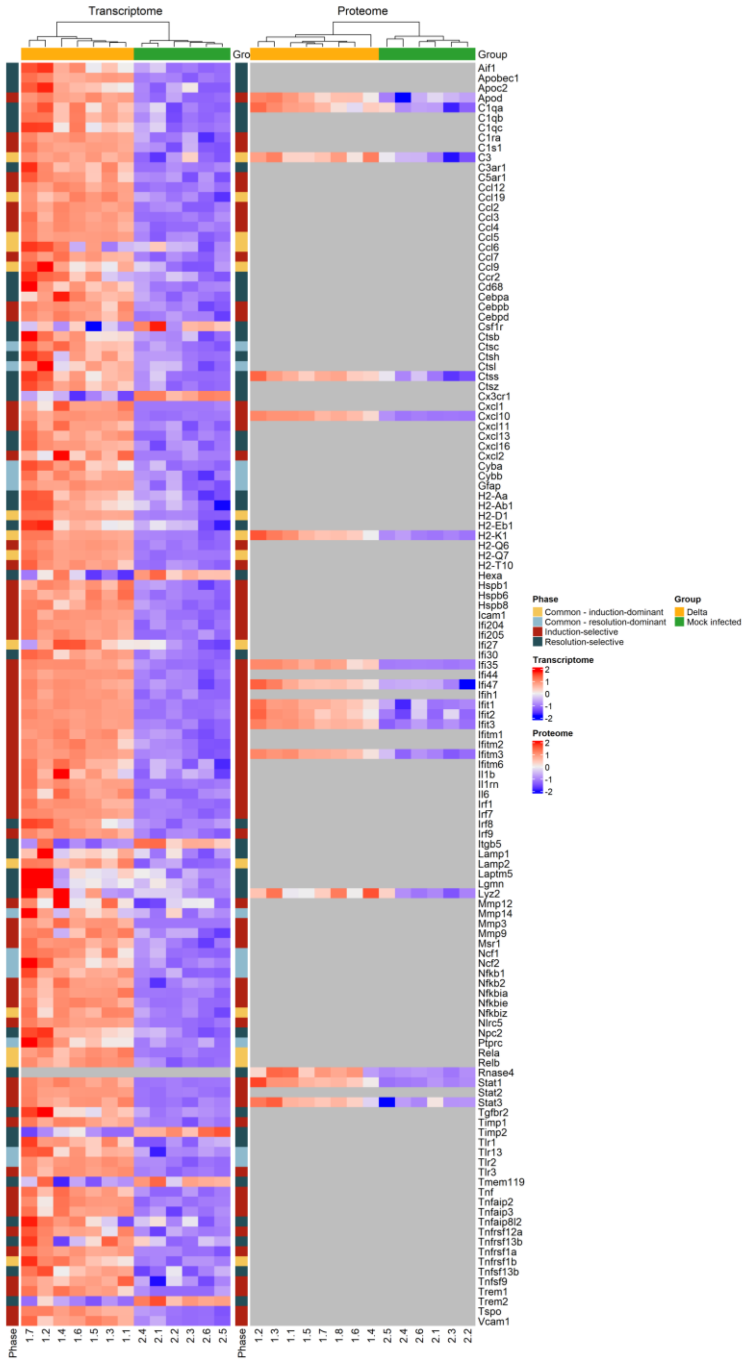


**Supplemental Figure S13. Transcriptome (thalamus) and proteome (cerebral cortex/hippocampus), K18-hACE2 mice after intranasal challenge with SARS-CoV-2 Delta variant ($10^2$ PFU/mouse) and mock-infected; 6-7 dpi.** Illustration of the significantly different gene products (False Discovery Rate < 0.05) derived from the brain inflammation signature from Davies and co-authors (89), in the Delta-infected (Delta) cohort and the mock-infected cohort.

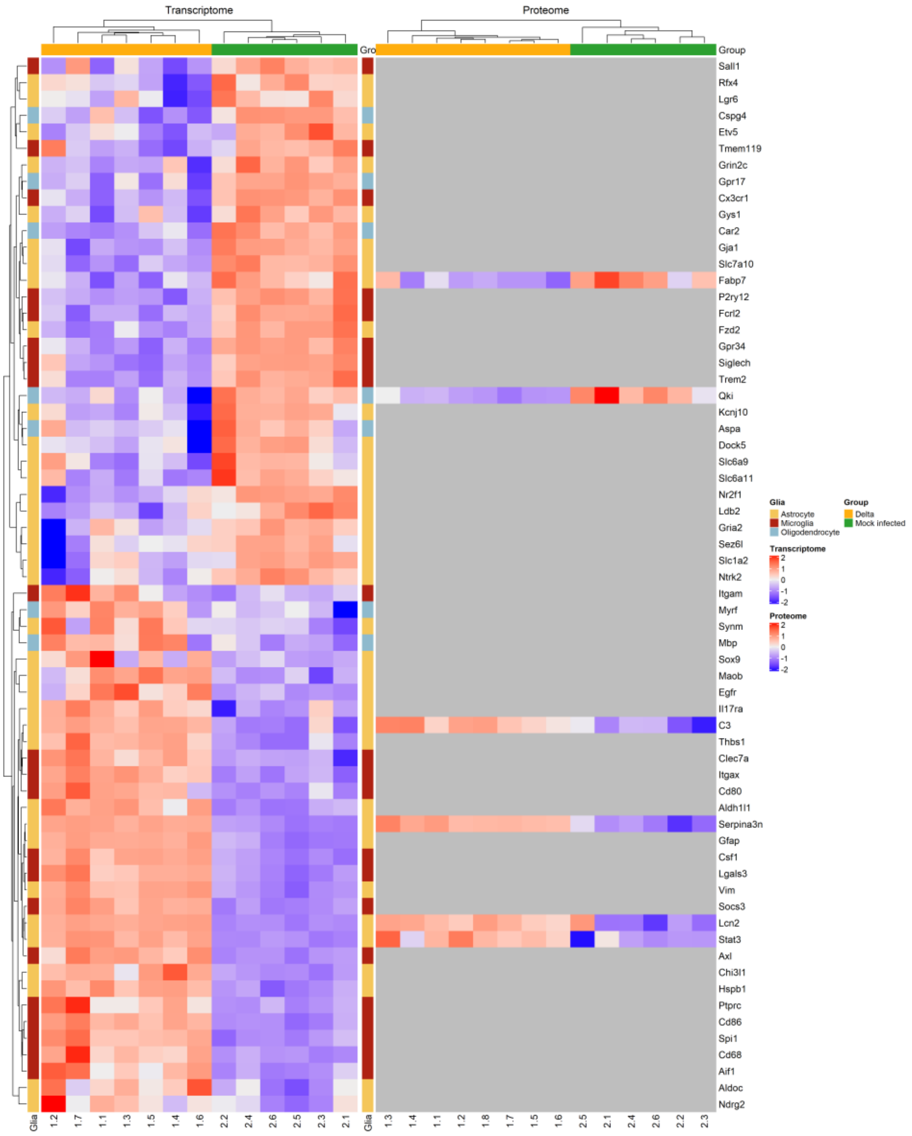


**Supplemental Figure S14. Transcriptome (thalamus) and proteome (cerebral cortex/hippocampus), K18-hACE2 mice after intranasal challenge with SARS-CoV-2 Delta variant ($10^2$ PFU/mouse) and mock-infected; 6-7 dpi.** Illustration of the significantly different gene products (False Discovery Rate < 0.05) between the Delta-infected (Delta) and the mock-infected cohorts, for several glial (astrocyte, microglia, oligodendrocyte) markers.

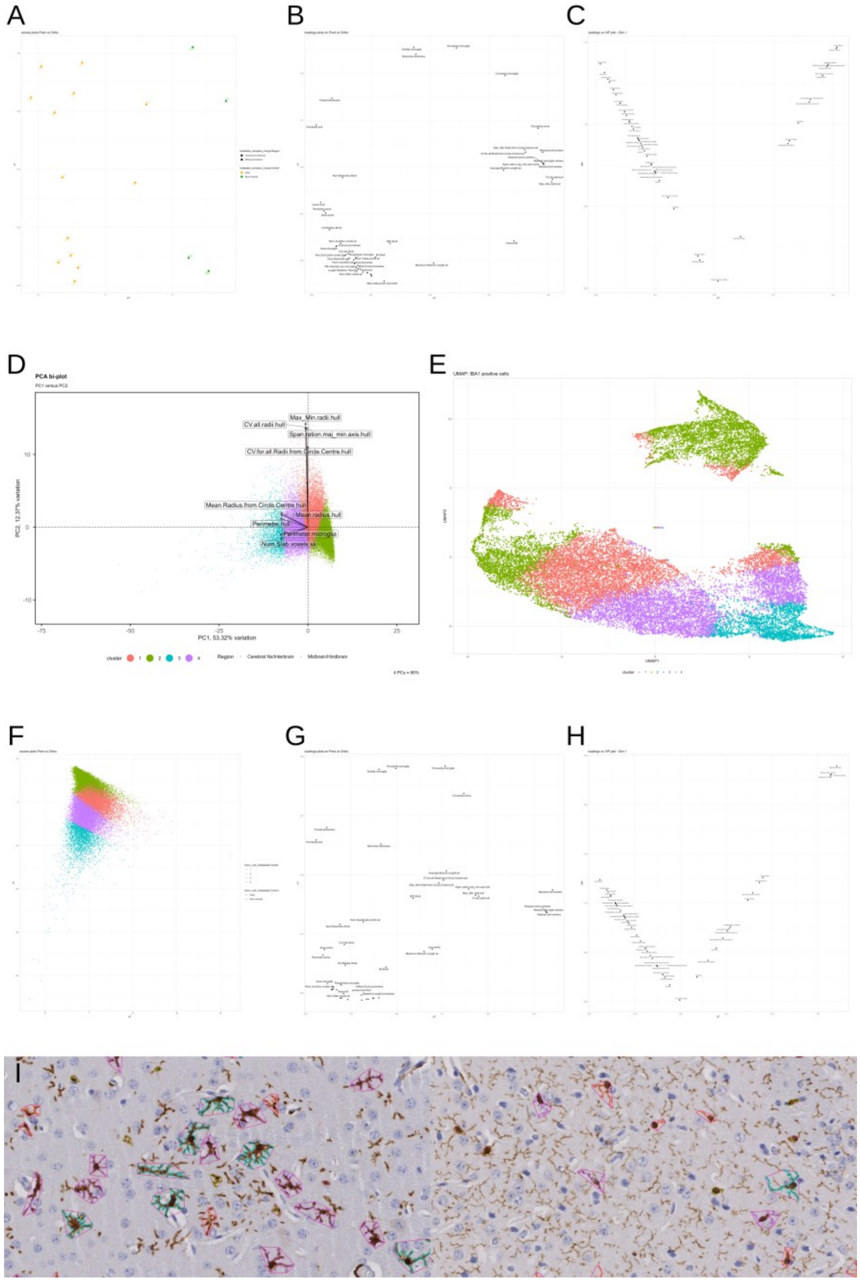


**Supplemental Figure S15. Brain, K18-hACE2 mice after intranasal challenge with SARS-CoV-2 Delta variant ($10^2$ PFU/mouse) and mock-infected; 6-7 dpi. Quantification of tissue expression of IBA1 and GFAP in the cerebral nuclei/interbrain and the midbrain/hindbrain**. **A-C)** Orthogonal Partial Least Squares Discriminant Analysis (O-PLS-DA) showing the clustering of the samples, using the median of the measurements derived from the QuPath, Skeleton, Fractal, and Sholl analyses, based on the cohort. **A)** Plot of the samples, showed a tendency to cluster within the respective groups (i.e., infected and mock-infected); **B)** Plot of the loadings; **C)** Plot of the Variable Importance in Projection (VIP). **D, E)** PCA (**D**) and Uniform Manifold Approximation and Projection (UMAP) (**E**) showing the clustering of the individual IBA1-positive cells. The shapes in the PCA biplot correspond to the brain region. **F-H)** O-PLS-DA showing the clustering of the individual IBA1-positive cells based on the cohort. **F)** Plot of the sample; **G)** Plot of the loadings; **H)** Plot of the VIP. **I)** Illustration of the IBA1-positive cells (i.e. microglia) associated to each cluster in the Delta-infected (left; mouse 1.1) and mock-infected (right; mouse 2.6) cohorts. Cerebral nuclei/interbrain. The colour coding of the clusters matches the colour coding in the PCA, UMAP, and O-PLS-DA plots.

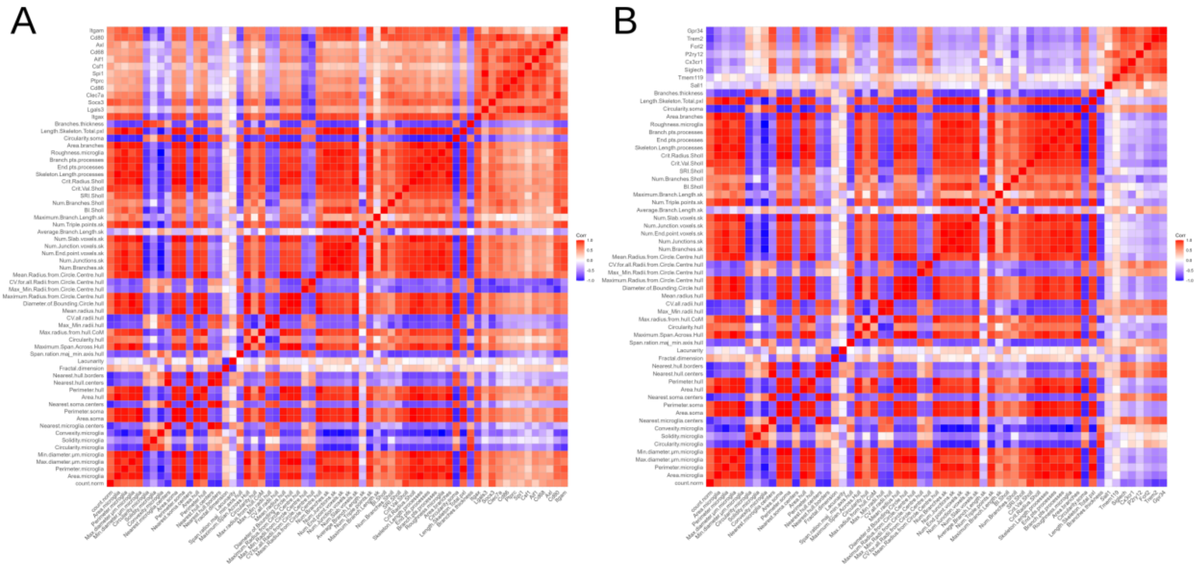


**Supplemental Figure S16. Brain, K18-hACE2 mice after intranasal challenge with SARS-CoV-2 Delta variant ($10^2$ PFU/mouse) and mock-infected; 6-7 dpi. Spearman correlation of the image analysis results at sample level (cerebral nuclei/interbrain) and the transcriptome (thalamus). A)** Correlation heatmap of the IBA1-positive cell measurements and the significantly up-regulated microglial markers (FDR < 0.05 and $log_2FC > 0$). **B)** Correlation heatmap of the IBA1-positive cell measurements and the significantly down-regulated microglial markers (FDR < 0.05 and $log_2FC < 0$). FDR: false discovery rate; $log_2FC$: $log_2$ fold change.

**Supplemental Material and Methods**

***Image analysis***

*Data analysis*

The results from the QuPath, Skeleton, Fractal, and Sholl analyses were imported in R (v.4.5.2, (90))/RStudio (RStudio: Integrated Development Environment for R, version 2026.01.0+392, Posit Software, PBC, Boston, Massachusetts, USA) for analysis. The R script detailing the analysis is available as a separate file (Supplemental MM-Digital pipeline step by step) and the list of R packages is available in Supplemental Table S8A. Briefly, upon import, the measurements from the four analyses were merged and each IBA1-positive cell was linked to its corresponding brain region (cerebral nuclei/interbrain or midbrain/hindbrain). IBA1-positive cells that could not be associated with a brain region were excluded from the analysis.

A first analysis focused on the comparison at sample level. For each sample, the median of each measurement, across all the IBA1-positive cells for that given sample and the corresponding brain region, were computed. Additionally, the number of IBA1-positive cells for each sample was counted and normalised to the area of tissue evaluated. The clustering of the samples was assessed using a Principal Component Analysis (PCA) (PCAtools, (45)), Orthogonal Partial Least Squares Discriminant Analysis (O-PLS-DA) (ropls, (47)), and Hierarchical Clustering Analysis (HCA) (cluster, (37)). The PCA and O-PLS-DA were performed on $log_2$-transformed, centered, and scaled data, while the HCA was performed on centered and scaled data. For all three, the measurement "Num.Quandruple.points.sk" was excluded as the value was "0" for each sample. For comparison between the groups, assessment of the normality was done with a Shapiro-Wilk test and statistical testing was done with the Mann-Whitney U test as not all measurements followed a normal distribution. The statistical analysis was done on each brain region separately (cerebral nuclei/interbrain and midbrain/hindbrain) and the alpha level was set at 0.05. As there were no significant differences between the groups, a similar analysis was done by taking all brain regions together. A correction for multiple testing was applied using the Benjamini & Hochberg method. The normalised counts of IBA1-positive cells for each sample, separated by brain regions, were visualised using barplots (ggplot2, (42)). Integration between the results of the IBA1 pipeline and the transcriptome (selecting for microglial markers; see Material and Methods "Transcriptome" for more details) was done via a correlation analysis (Spearman). For the correlation, the measurements from the cerebral nuclei/interbrain were used, as the transcriptome analysis was done on the thalamus.

As the histological investigation revealed clear differences in the dominant microglia shape between the infected and control groups, we investigated the individual IBA1-positive cells to determine if the measurements obtained from the digital pipeline would also indicate a difference. To do so, a k-means clustering analysis was performed (MacQueen algorithm using four clusters) and the results were investigated with a PCA, O-PLS-DA, Uniform Manifold Approximation and Projection (UMAP) (umap, (53)), and T-Distributed Stochastic Neighbor Embedding (t-SNE) (Rtsne, (49)). Additionally, the results were exported back to QuPath, and each IBA1-positive cell was colour-coded based on its cluster, allowing for visual assessment and quality control. Of note, for the clustering analysis, the individual cells which had a NA value in SRI.sholl and/or Branches.thickness measurements were excluded. Because these two parameters evaluate the branching, and we have histologically determined that the microglia in both groups have clearly identifiable branches, excluding these cells is considered not to significantly impact the overall conclusion. Additionally, in an attempt to improve the clustering, an analysis was performed using only the parameters which were significantly different between the two groups. However, because of the suboptimal results and separation between the clusters, this analysis was not pursued further.

For both approaches (samples or individual cells) the performance of the models was evaluated with a permutation test (random permutations, n = 100); the quality metrics of the models are summarised in the table below.

| Mode | R2X (cum) | R2Y (cum) | Q2 (cum) | RMSEE | pR2Y | pQ2 |
|---|---|---|---|---|---|---|
| Sample | 0.867 | 0.789 | 0.698 | 0.215 | 0.01 | 0.01 |
| Cell | 0.623 | 0.209 | 0.209 | 0.279 | 0.01 | 0.01 |

*Limitations*

The pipeline used for the automated image analysis of the IBA1-positive cells is associated with some inherent limitations. To select complete microglial cells (i.e., the soma and cell processes), we had to exclude smaller IBA1 detections, which excluded many cell processes in the mock-infected cohort and biased the representation of a "normal" microglial cell as seen in a control brain. Additionally, the lower number of IBA1-positive detections in the control brains means the dataset used for clustering was biased toward microglia with altered morphology. Also, because IBA1 expression does not allow to distinguish microglial cells from monocytes/macrophages, some detections in the infected brains will have corresponded to individual or overlapping monocytes/macrophages.

***3D fluorescence imaging***

Volumetric image of SARS-CoV-2 distribution across the entire brain was provided as a service by Clepio Biotech Srl (Florence, Italy, www.clepiobiotech.com). For this, the brain of a mouse from a separate experiment was used (SARS-CoV-2 Delta at a dose of $10^3$ PFU/mouse, 8 days post infection (dpi)). Briefly, after fixation with paraformaldehyde, the entire brain was cleared and stained according to the iDISCO protocol (91). Immunostaining against SARS-CoV-2 nucleocapsid protein (NP) was performed with the anti-SARS-CoV-2 NP antibody used in immunohistochemistry and immune electron microscopy (see Material and Methods (Histological and immunohistological examinations) in the main text) at a dilution of 1:2500, followed by a secondary antibody stained with an Alexa 647 dye. After staining and clearing, samples were imaged with high-resolution light-sheet fluorescence microscopy (92). Raw imaging data were stitched, then compressed and saved using a multi-resolution Zarr file format (93). The final data was visualised using napari (v0.10.0 (94)).

***Omics analysis***

**Transcriptome (single -omic)**

For the cell deconvolution, the reference matrix of murine thalamus was built using the data from Zeisel et al. (95). To this end, the "l5_all.agg.loom" file, containing the non-normalised averaged UMI counts per cell type, was downloaded from the companion wiki (http://mousebrain.org/adolescent/downloads.html, accessed on the 12-05-2025) (at the time of writing, the above link was not accessible anymore; the paper now redirects to the following companion wiki https://linnarssonlab.org/mousebrain/). Upon import in R/RStudio, the file was filtered to keep only the cell types associated with the thalamus, central nervous system (CNS), and brain. CIBERSORTX (96) was run in HiRes mode, with rmbatchBmode set to TRUE, to compute the cell fractions. The signature matrix was the filtered file from Zeisel et al. (95) and the mixture file was our non-normalised and non-filtered transcriptome dataset. Additionally, a classes file was provided in order to group cell types in broader categories (neuroblast-like, inhibitory neuron, cholinergic neuron, excitatory neuron, oligodendrocyte, choroid plexus, ependymal, astrocyte, vascular leptomeningeal cell, endothelial cell, pericyte, smooth muscle cell, perivascular macrophage, microglia) (Supplemental Table S9); this was done using the companion wiki (http://mousebrain.org/adolescent/celltypes.html, accessed on the 15-05-2025). The results were visualised with a barplot (ggplot2, v4.0.1, (42)) and the cell fractions were automatically adjusted so that their total, per sample, was 1. For inhibitory neuron, excitatory neuron, astrocyte, and microglia, following a Shapiro-Wilk test, and a F-test when appropriate (inhibitory neuron, excitatory neuron, astrocyte), statistical significance between the infected cohort and the mock-infected cohort were assessed with either a t-test (inhibitory neuron, excitatory neuron, astrocyte), or a Mann-Whitney U test (microglia). In all tests, the alpha level was set at 0.05.

**Proteome (single -omic)**

While in our previous publication (14) we focused only on the host proteome, in this study we also investigated the SARS-CoV-2 proteome. While both the host and SARS-CoV-2 proteins were acquired in the same batch, we adapted the post-acquisition pipeline to minimise artefacts for each proteome. As such, to minimise quantification artefacts arising from match-between-runs (MBR)-mediated transfer of SARS-CoV-2 protein identifications from infected to mock-infected samples, viral protein abundances were determined using data processed with MBR disabled. On the other hand, for the host proteome, we used data processed with MBR enabled, similarly as in our previous publication (14). Upon import of the two datasets in R/Rstudio, each one underwent a specific pipeline, focusing either on SARS-CoV-2 proteins (MBR disabled), or on host proteins (MBR enabled), using for each the MaxLFQ intensity as the protein abundances.

For the SARS-CoV-2 proteome pipeline, upon import in R/RStudio, the protein abundances were normalised using variance stabilizing transformation (vsn) (using DEP2, (62)), applied on both the host and SARS-CoV-2 proteins, after which SARS-CoV-2 proteins were selected for statistical analysis and visualisation. The normalised SARS-CoV-2 protein abundances were visualised with a heatmap (using pheatmap, (77)) and boxplots (ggplot2, (42)). Following the assessment for normality with a Shapiro-Wilk test, comparison between the Delta- and mock-infected cohorts, for each SARS-CoV-2 protein, was performed with a Mann-Whitney U test. In all tests, the alpha level was set at 0.05. Details on the R packages are available in Supplemental Table S8.

The host proteome pipeline is described below, in the “Transcriptome-proteome integration (high-level integration)” section.

**Metabolome (single -omic)**

Functional enrichment analysis was performed using the FELLA package (66), following the authors’ instructions, with KEGG (97–100). After building the KEGG graph (done on the 27.01.2026), the enrichment analysis was performed using the list of metabolites which were significantly different between the Delta-infected and the mock-infected cohorts (FDR < 0.05) and for which a KEGG compound ID (cpd) was available (mapping done with RefMet (80)). The input metabolites were: citraconic acid, quinic acid, uric acid, S-adenosylhomocysteine, methyl beta-D-galactoside, and pantolactone. For the enrichment analysis, the method argument was set to “diffusion” and the approx argument was set to “normality”. The metabolites methyl beta-D-galactoside and pantolactone could not be mapped to the built KEGG graph. The subgraph with the mapped metabolites, and the result tables, were exported for data exploration.

The abundance of the neurotransmitters glutamate and GABA were visualised with violin plots (ggplot2, (42)).

Details on the R packages are available in (14) and in Supplemental Table S8B.

**Lipidome (single -omic)**

As the PCA did not show separation of the two cohorts (infected, mock-infected) (14), we attempted to cluster the data using an O-PLS-DA to further investigate potential lipids of interest associated with the brain infection. The O-PLS-DA (ropls, (47)) was done on the filtered, median normalised, and $log_2$-transformed data set (14); the separation was based on the cohort (infected, mock-infected). The lipidome positive and negative mode data sets were processed separately. The performance of the models was evaluated with a permutation test (random permutations, n = 100); the quality metrics of the models are summarised in the table below.

| **Mode** | **R2X (cum)** | **R2Y (cum)** | **Q2 (cum)** | **RMSEE** | **pR2Y** | **pQ2** |
|---|---|---|---|---|---|---|
| Positive | 0.353 | 0.923 | 0.69 | 0.155 | 0.06 | 0.02 |
| Negative | 0.367 | 0.874 | 0.473 | 0.198 | 0.05 | 0.06 |

Functional enrichment analysis was performed on the positive mode datasets, as no significant lipids were detected in the negative mode, using the LSEA method within LipidSigR (72) and the LION ontology (101, 102). The abundance of the lipids (positive mode) was normalised (median), $log_2$-transformed, centred and scaled, and the TG and TG O- lipid species were visualised with a heatmap (pheatmap, (77)).

Details on the R packages are available in (14) and in Supplemental Table S8B.

**Multi-omics: Low-level integration**

The low-level multi-omics integration was based on a multiblock analysis approach, using either an unsupervised (Common Dimension method (ComDim) (75)) or a supervised (consensus orthogonal PLS-DA (O-PLS-DA) (60, 103)) method. For each method, each -omic dataset (i.e. transcriptome (n = 16388), proteome (host: n = 5289), metabolome (n = 130), lipidome (positive mode) (n = 784), lipidome (negative mode) (n = 27)) was derived from the corresponding single -omic pipeline. These datasets were filtered, normalised, $log_2$-transformed, scaled and centered, individually, before performing the multiblock analysis. The ComDim method was performed using the ComDim() function from the MBAnalysis package (75), and scaling between blocks was computed via the scale.block argument. The results were visualised with block contributions, score, and loading plots, using ggplo2 (42). The consensus O-PLS-DA was computed using the cohort (infected, mock-infected) as the explanatory variable. The number of predictive and orthogonal components were each set to 1 and the kernel was set to polynomial. Performance of the model was assessed via cross-validation (Leave-One-Out cross validation) using 100 random permutations. The results were visualised with block contributions, score and loading plots as well as variable influence on projection (VIP) versus loadings on predictive component, using ggplo2 (42). The R-squared ($R^2Y$), Stone Geisser $Q^2$ (Q2), and discriminant $Q^2$ (DQ2) index are summarised in the table below, and the distribution of the permuted values is illustrated in Supplemental Figure S10. Details about the R package used in the analysis are available in Supplemental Table S8B.

| R2Y | Q2 | DQ2 |
|---|---|---|
| 0.9984 | 0.8682 | 0.8467 |

**Multi-omics: high-level integration**

***Transcriptome-proteome integration***

The transcriptome and proteome were analysed in parallel and integrated, with a focus on SARS-CoV-2 cycle/replication, innate immunity/antiviral response, glial response, and neurons.

Results from the functional enrichment analysis from both the transcriptomic and proteomic analyses (Gene Ontology (104, 105) and Reactome (106, 107) databases, using GSEA via clusterProfiler (55–58)), were manually reviewed to select terms of interest. The results from the two -omics analyses were visualised together with dotplots (ggplot2, (42)).

List of genes of interest were derived from the literature for the following categories: 1) interferon-stimulated genes reported in SARS-CoV-2 infection (108–121); 2) innate immunity/antiviral response (120, 122–131); 3) Pattern Recognition Receptors (PRRs) (132, 133); 4) glial response (95, 134–165); 5) signature of the resolution of inflammation (89); 6) neuronal markers (137, 165–176); 7) neuropeptides, neurotransmitters, synapses (165, 176, 177). Using these gene lists, the differentially expressed gene products were parsed, and the corresponding $\log_2$-transformed and row scaled counts and intensities were visualised using heatmaps (ComplexHeatmap, (59)) for gene products of interest. For the "neuronal markers" and the "neuropeptides, neurotransmitters, synapses categories", to allow for comparison between the abundance of the gene products in the transcriptome and proteome, the significant gene products (false discovery rate < 0.05) were mapped to the significant, or non-significant, gene products in the proteome and transcriptome respectively.

A list of the packages used is available in Supplemental Table S8B.
